# Optimized Quantification of Intrahost Viral Diversity in SARS-CoV-2 and Influenza Virus Sequence Data

**DOI:** 10.1101/2021.05.05.442873

**Authors:** AE Roder, KEE Johnson, M Knoll, M Khalfan, B Wang, S Schultz-Cherry, S Banakis, A Kreitman, C Mederos, J-H Youn, R Mercado, W Wang, D Ruchnewitz, MI Samanovic, MJ Mulligan, M Lassig, M Łuksza, S Das, D Gresham, E Ghedin

## Abstract

High error rates of viral RNA-dependent RNA polymerases lead to diverse intra-host viral populations during infection. Errors made during replication that are not strongly deleterious to the virus can lead to the generation of minority variants. However, accurate detection of minority variants in viral sequence data is complicated by errors introduced during sample preparation and data analysis. We used synthetic RNA controls and simulated data to test seven variant calling tools across a range of allele frequencies and simulated coverages. We show that choice of variant caller, and use of replicate sequencing have the most significant impact on single nucleotide variant (SNV) discovery and demonstrate how both allele frequency and coverage thresholds impact both false discovery and false negative rates. We use these parameters to find minority variants in sequencing data from SARS-CoV-2 clinical specimens and provide guidance for studies of intrahost viral diversity using either single replicate data or data from technical replicates. Our study provides a framework for rigorous assessment of technical factors that impact SNV identification in viral samples and establishes heuristics that will inform and improve future studies of intrahost variation, viral diversity, and viral evolution.

**IMPORTANCE:** When viruses replicate inside a host, the virus replication machinery makes mistakes. Over time, these mistakes create mutations that result in a diverse population of viruses inside the host. Mutations that are neither lethal to the virus, nor strongly beneficial, can lead to minority variants that are minor members of the virus population. However, preparing samples for sequencing can also introduce errors that resemble minority variants, resulting in inclusion of false positive data if not filtered correctly. In this study, we aimed to determine the best methods for identification and quantification of these minority variants by testing the performance of seven commonly used variant calling tools. We used simulated and synthetic data to test their performance against a true set of variants, and then used these studies to inform variant identification in data from clinical SARS-CoV-2 clinical specimens. Together, analyses of our data provide extensive guidance for future studies of viral diversity and evolution.

## INTRODUCTION

Large population sizes, high replication rates, and error prone polymerases all contribute to the generation of sequence diversity found in viral infections (1–5). Natural selection acts on this diversity, contributing to viral evolution. RNA viruses have some of the highest mutation rates among viruses (1, 6, 7). To replicate their genomes, RNA viruses must encode their own RNA-dependent RNA-polymerases (RdRp), which often lack proofreading capabilities. Coronaviruses are a notable exception, as they possess a distinct protein with 3’-5’ exonuclease capability (1, 8, 9). Most errors made during replication—up to 40% in RNA viruses—are lethal (10, 11). Beneficial mutations make up a much smaller proportion, and these, along with neutral mutations, comprise the substitution rate. This substitution rate can be used to estimate the viral evolutionary rate, an important calculation in considering viral spread, pandemic potential, and vaccine design (4, 12). Due to the large population sizes of RNA viruses, intrahost bottlenecks, and genetic drift, genetic diversity within a host is dynamic, with frequencies of mutations constantly rising and falling (13). Mutations can lead to changes in the consensus sequence, e.g., where the allele frequency (AF) is greater than 50%, and these specific sets of mutations separate globally circulating virus populations into clades. Mutations in the virus genomes that are not the majority within an infected host (i.e., present at lower than 50% frequency) represent minority variants. Deep sequencing enables the capture of intrahost variation, both at the majority and minority level, enabling the identification of variants and estimation of their frequency. Studying intrahost variation can help in tracking viral spread, estimating population bottleneck sizes, and identifying key amino acid changes that differentiate new viral strains (14–17). Additionally, minority variants can highlight regions of the genome under selection or regions with increased mutational tolerance, as well as allow for detection of subtle population shifts within the infected host and discovery of possible drug resistance mutations (18, 19). Thus, information gleaned from studying intrahost viral diversity has major implications for vaccine, monoclonal antibody, and drug development.

Given the many applications of studying intra-host viral diversity, accurately identifying and quantifying viral variants is essential. Precise identification of viral variants, especially those at low frequencies, is complicated by the fact that viral genome sequencing often requires reverse transcription and amplification, which, along with library preparation and the sequencing process, are error prone. Thus, distinguishing true sequence variation from technical and experimental noise is challenging. Typically, several ad hoc metrics are used to filter variants, such as applying frequency and coverage cutoffs to sequencing data, however, the frequency at which identified variants are considered valid can vary widely (20–27). Most studies using large sample cohorts, or performing analyses on publicly available data, generally use single replicate data, despite evidence suggesting that replicate sequencing may be essential for filtering false positive minority variants (27). Despite the large number of studies analyzing minority variants in virus data, there is no consensus on what coverage cutoffs and allele frequency cutoffs to use, and no large-scale studies have been performed to determine what thresholds lead to the highest confidence in variant identification.

In addition to the diversity of cutoffs used for single nucleotide variant (SNV) identification, there is also great diversity in the variant calling software available. Variant callers are often designed with specific functions in mind, such as identifying germline or somatic mutations in cancer genomes or single nucleotide variants in viral populations (28, 29). The function for which a variant caller is designed can have a significant impact on the statistics used and assumptions made by the software. Tools designed for detection of germline mutations, such as HaplotypeCaller and freebayes, must consider the very large reference genome, higher frequency variants, the diploid nature of the genome, the possibility of copy number variation, and long repetitive regions or large insertions or deletions (30–35). In these instances, local realignment of haplotypes may be most effective (28). By contrast, software used for somatic mutations in tumors, such as Mutect2 and Varscan, or for viral diversity, such as iVar and timo (a variant caller developed in our lab), use base by base comparisons, or a combination of this with haplotype-based alignment, to find lower frequency variants (27, 33–36). These tools also may need alternative methods to reduce false positive calls to account for PCR errors introduced during amplification of the viral genome (29). Due to the differences in bioinformatic and statistical approaches used by each variant calling tool, identifying the tool that is the best fit for the specific research question being studied is essential. Some tools have been tested in pairwise comparisons (27, 35), however little work has been done to extensively test the performance of the many available tools on different viruses, across sequence coverages, and at various allele frequencies in viral deep sequencing data.

Here, we tested seven variant callers on simulated, synthetic, and clinical deep sequencing data. We tested each tool across a range of coverages, allele frequencies, and experimental designs to determine the optimal parameters that should be used to decrease false positive variant identification, without sacrificing true positive data. To compare performance between a small RNA virus with a high mutation rate, and a large RNA virus with proofreading capability, we tested the variant callers on two viruses of particular interest in the viral diversity field, influenza and SARS-CoV-2. We find that choice of variant caller, and use of replicate sequencing have the most significant impact on SNV discovery and demonstrate how both allele frequency and coverage thresholds impact both false discovery and false negative rates. We also provide guidance on best practices for leveraging deep sequencing data from public repositories for intrahost studies. These analyses provide a resource for studies aiming to assess intrahost viral diversity in SARS-CoV-2 or influenza, and they lay the groundwork for similar studies in other viruses.

## MATERIALS AND METHODS

### Extended methods are available in the supplementary materials

#### Generation of Simulated Data

Reads were simulated using NEAT (v2.0) by constructing a mutation, error, fragment length, and GC model for each viral type (37). The models were provided to NEAT genReads.py along with reference fasta files and a mutation rate of 0.009 (0.9%) for influenza and 0.0045 (0.45%) for SARS-CoV-2 to produce a “golden VCF” containing a defined number of SNVs in each virus. Simulated random PCR errors were also added to each replicate using genReads.py (NEAT). Several copies of the replicate golden VCFs were made, each with the same variants but with differing allele frequencies (AF). These VCFs were used to simulate paired end fastq libraries at 100,000X genome coverage and down-sampling was used to simulate lower coverages.

Sequences were trimmed using trimmomatic v0.36 (38), aligned to the respective reference genome with BWA mem v0.7.17 (39), and duplicate reads were marked using GATK MarkDuplicatesSpark v4.1.7.0 (40). Variants were called in each replicate with seven different tools, using multiple parameter configurations for each tool (**Table S1)**. A VCF file containing the intersection of the two replicates was generated using bcftools isec (v1.9) (39). The pipeline used for data simulation, sequence processing, variant calling, and analysis is available at https://github.com/gencorefacility/MAD2.

##### Synthetic RNA generation, library preparation, and data processing

Synthetic, ‘wildtype’ (WT), influenza genomic RNA and variant RNA (created by adding 18, 14 and 14 known nucleotide changes into the WT PB2, HA, and NA segments respectively) were synthesized as double-stranded DNA (gBlocks). The DNA was *in vitro*-transcribed with the HiScribeTM T7 High Yield RNA Synthesis Kit (Invitrogen). RNA samples were diluted to an equal copy number concentration of 6×10^8^ copies/µL. WT and variant RNA were mixed at equal molarity. The pools were mixed at various frequencies (50%, 25%, 12.5%, 6.25%, 3.13%, 1.56%, 0.78%, 0.39%), and diluted to various copy number concentrations (6×10^6^ - 6×10^3^ copies/µL).

cDNA was generated and libraries were prepared using the Nextera XT library preparation kit (Nextera), with all volumes scaled down to 0.25x of the manufacturer’s instructions, cleaned with AMPure beads, and pooled at equal molarity. Libraries were sequenced on the Miseq 300 Cycle v2 using 2 × 75 pair-end reads. Samples were amplified and sequenced in duplicate and analyzed with the pipeline described above, with the addition of adapter trimming. Synthetic SARS-CoV-2 data from a similarly designed study was downloaded from SRA (PRJNA682212) and processed as above (26).

### SARS-CoV-2 clinical sample preparation, processing, and variant calling

Total RNA was extracted from 300 µL of nasopharyngeal (NP) or mid-turbinate (MT) swabs collected at the NIH Clinical Center as part of diagnostic testing between 07/24/2020 and 03/31/2021 (**Table S2**). All samples were de-identified and anonymized.

RNA from samples was extracted using the NUCLISENS easyMAG automated nucleic acid extractor and the viral genome was amplified using a modified version of the ARTIC protocol (https://artic.network/ncov-2019) and the methods described at https://github.com/GhedinSGS/SARS-CoV-2_analysis. All libraries were prepared as above and sequenced on either the Illumina MiSeq or the Illumnia NextSeq500 using either the 2×150 bp or 2×300 bp paired end protocol. All samples were processed in duplicate.

Samples were processed with the pipeline available and described above, with the addition of merging the two SAM files (from A and B primer pools) for each biological sample into one alignment file using Picard Tools MergeSamFiles v2.17.11. Variants were called as above using the standard parameters for each tool (**Table S1**).

### Data Availability

Synthetic influenza data (bioproject PRJNA865369) and SARS-CoV-2 data from clinical samples are available in NCBI GenBank and SRA. Accession IDs can be found in **Table S2**. All downstream analysis files are available at https://github.com/GhedinSGS/Optimized-Quantification-of-Intrahost-Viral-Diversity.

### Ethics Statement

All samples were anonymized and obtained with consent as part of SARS-CoV-2 diagnostic testing.

## RESULTS

### Simulated and synthetic data provide a ‘true’ set of minority variants to assess variant caller performance

To test the ability of each variant caller to accurately identify variants, it is essential to know the ‘true set’ of variants within the data, which is not possible with real sequence data. With this in mind, we tested the ability of six popular variant-calling software packages (Freebayes, HaplotypeCaller (hc), iVar, Lofreq, Mutect2, and Varscan) and one in-house pipeline (timo) to accurately identify minority variants in simulated and synthetic sequencing data (**Fig. S1**) (19–24). Single nucleotide variants (SNVs) were simulated across three influenza virus genomes (A/H1N1, A/H3N2, B/Victoria) and one coronavirus genome (SARS-CoV-2) at both defined and random allele frequencies and across a range of downsampled coverages (**Fig. S1A-B**). Further, synthetic RNA controls of three influenza virus segments (PB2, HA, and NA) containing known SNVs were mixed in varying amounts at various dilutions to create a range of allele frequencies and genome copy numbers (**Methods**) (**Fig. S1C-D**). Combined, we used these synthetic and simulated data sets to test variant caller performance on a known set of SNVs.

We found that all callers performed poorly on our data using their default parameters (**Fig. S2A**). Therefore, to compare all callers equally, we used a standard set of permissive input parameters (min coverage = 1x, allele frequency cutoff = 0.01 (1%)) throughout our testing (**Table S1**). When assessing the F1 statistic across a range of simulated frequencies, most variant callers performed well at low frequencies (< 0.05 (5%)) when the coverage was high (downsampling fraction >= 0.005 or ∼500X coverage). Conversely, high frequencies were necessary for accurate variant detection at small downsampling fractions where the average coverage was low (**Fig. 1A**). We did not observe significant differences between the four viruses in terms of variant calling accuracy (**Fig. 1A, Fig. S2A**). Most of the differences in performance between the variant callers could be seen at downsampling fractions < 0.005 and allele frequencies below 0.05 (5%) (**Fig 1A**). A closer look at precision and recall for each tool at downsampling fractions of 0.002 and 0.003 indicated that many tools trade recall for precision at low frequencies (**Fig. 1B**). Some tools, such as iVar, timo, and varscan tended to be extremely conservative, especially at low frequencies. However, the stringency of these tools can be reduced by decreasing the input frequency parameter from 0.01 (1%) to 0.001 (0.1%) (**Fig. S2** *custom input parameters*, **Table S1**). The use of simulated data for initial testing showed that under ideal conditions, all variant callers have the ability to perform well on viral sequence data. This is important for establishing baseline performance against which to compare the variant caller performance on synthetic data and data from clinical specimens.

**Figure 1.**
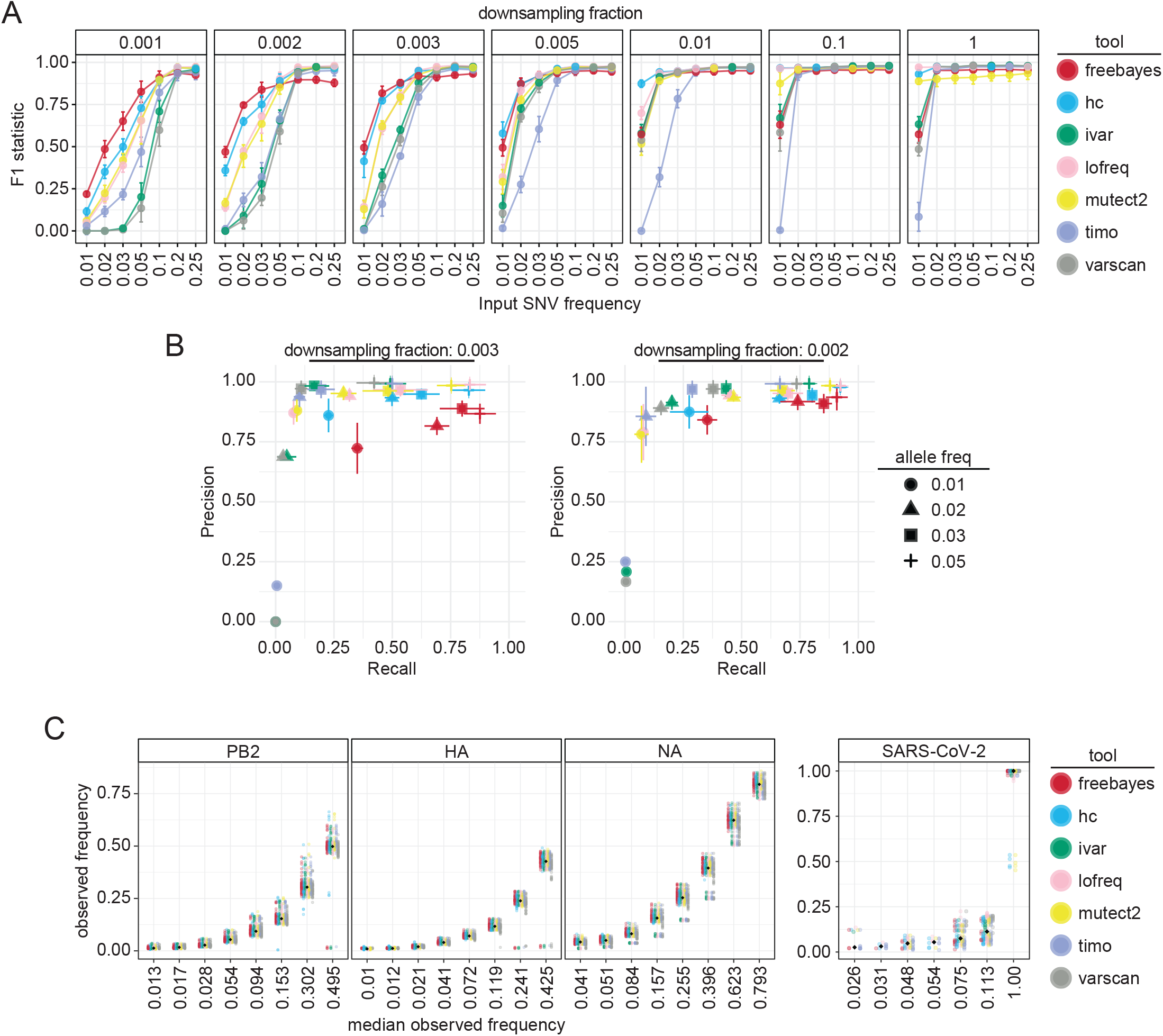
Variant caller performance on simulated and synthetic data with variants at set allele frequencies. **(A)** F1 statistic for each variant caller across a range of downsampling fractions and allele frequency values. Values shown are mean and standard deviation of the four viruses (A/H1N1, A/H3N2, B/Victoria, SARS-CoV-2) using standard input parameters (**Table S1**). Color represents the variant caller used. **(B)** Precision/Recall graphs of each variant caller across allele frequencies (point shape) for downsampling fractions 0.003 (top left) and 0.002 (top right) or across downsampling fractions (point shape) for allele frequencies (AF) 0.02 (bottom left) and 0.03 (bottom right). Color represents variant caller. Mean and standard deviation are shown across the four viruses for precision and recall scores. **(C)** Scatter plot showing median observed frequency (x-axis) versus individual observed frequency (y-axis) for each synthetic influenza gene segment and SARS-CoV-2 genome. Black dots indicate the median value for each expected frequency. Color represents the variant caller used.

While simulated data provides an ideal set of variants against which to compare the variant calls for each tool, it lacks the reverse transcription, amplification, and sequence library preparation steps involved in the generation of data from clinical specimens. To assess how these sample preparation steps, along with duplicate sequencing, and SNV thresholds may impact variant caller performance, we tested each tool on data from the synthetic RNA dataset (**Fig. S1C-D**). The average read depth across gene segments was greater than 1000x and had similar coverage distributions to our simulated datasets at downsampling fractions of 0.01 and 0.1 (**Fig. S3A**). At this coverage in simulated data, we observed high F1, precision, and recall scores across all variant callers for SNVs > 1% frequency (**Fig. 1A-B**). Observed frequencies differed from expectation (**Fig. S3B**), likely due to mixing errors during sample preparation, but were consistent between viral copy numbers. Comparing the observed allele frequency to the median observed allele frequency revealed that most callers agreed on the frequency of identified variants; however, there is considerable variance in the AF estimations despite the fact that all SNVs are linked (**Fig. 1C, S3B**). We also tested the variant callers on a previously published synthetic SARS-CoV-2 data set (26). Here, HaplotypeCaller and timo found more true variants than the other callers, especially at higher viral load (**Fig. S3C**). Across all callers, the frequencies of observed variants in the SARS-CoV-2 data were less accurate than in the synthetic flu data i.e. the variance in allele frequencies is higher) (**Fig. S3B-C**). This is potentially due to the overall low coverage of these samples and the amplicon-based sequencing method required for larger RNA genomes, suggesting that allele frequency estimation may be less reliable in SARS-CoV-2 data.

### Frequency thresholds and sequencing replicates reduce false positive SNVs

Previous studies have reported the necessity of establishing frequency and coverage thresholds as well as having replicate sequencing to decrease false-positive SNVs in a data set (27, 40). Given that most publicly available data consist of single replicate sequencing data, we aimed to establish coverage and frequency thresholds that would minimize the false discovery rate (FDR) and false negative rate (FNR) to levels comparable to those observed using two replicates. To do this, we used both simulated and synthetic datasets with standard input parameters (**Fig. S2A**) and ignored the ‘binocheck’ requirement from timo (requiring variants to be found in near equal numbers of forward and reverse reads), allowing us to test the performance of timo on low frequency SNVs.

In both the synthetic and simulated data, false positive SNVs were found across read depths but were primarily at allele frequencies less than 0.03 (**Fig. 2A-2B**). Therefore, applying frequency thresholds to single replicate data lowered the false discovery rate (FDR) for all callers (**Fig. 3A, S4A**). While establishing coverage cutoffs did not drastically impact the number of false positive calls in either the simulated or synthetic dataset, coverage and library size are important factors when considering SNV recall (**Fig. 1A, 2A-B, S5**). However, using frequency thresholds did come at the cost of significantly increasing the FNR, as true SNVs found at low frequencies were filtered from the data. In contrast, keeping only SNVs shared between the two replicates dramatically decreased the FDR, while maintaining relatively low FNRs (**Fig. 3, S4**). HaplotypeCaller, LoFreq, and Mutect2 called notably higher numbers of false positive SNVs in synthetic data, including many that were maintained even after merging replicates—indicating that these callers are making consistently incorrect SNV calls. Further, these three callers had multiple instances where true positive SNVs were identified at high frequencies (AF > 0.05) in one replicate and were entirely absent in the other (**Fig. S6A**). Importantly, replicates also increased the accuracy of allele frequency estimation of true positive variants in simulated data. The effect of sequencing replicates on allele frequency is especially pronounced for low coverage data, where the percent error of allele frequency estimation is pointedly lower for all tools when using replicates (**Fig. S6B**). This effect was less pronounced in synthetic data, likely due to the high coverage of these samples, again demonstrating the importance of read coverage for accurate allele frequency estimation (**Fig. S6C**).

**Figure 2.**
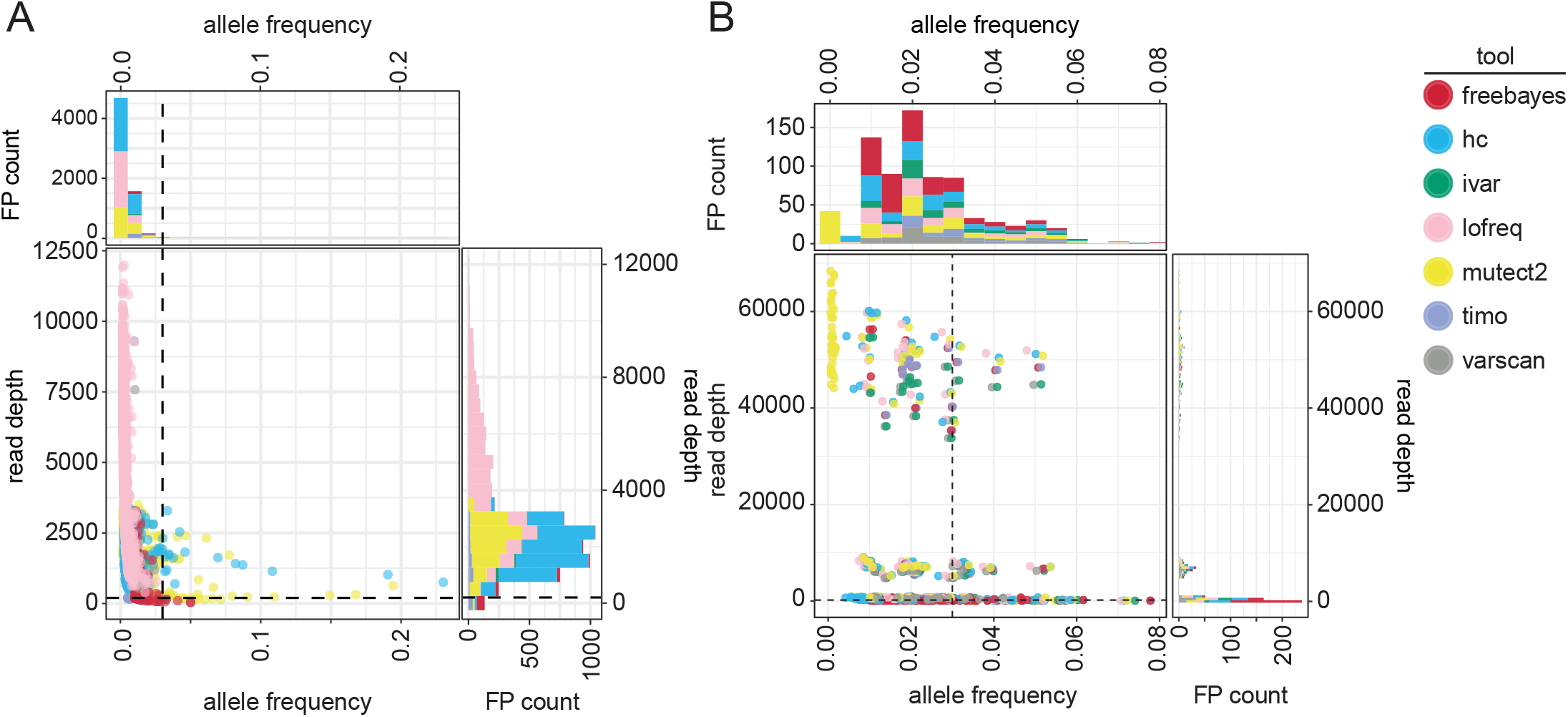
Frequency and coverage of false positive variants in synthetic and simulated data. **(A-B)** Scatter plots and associated histograms showing number of false positive SNVs across allele frequencies and read depths in synthetic (**A**) or simulated (**B**) data. Dotted lines are drawn at allele frequency = 0.03 and read depth = 200x. Color represents the variant caller used.

**Figure 3.**
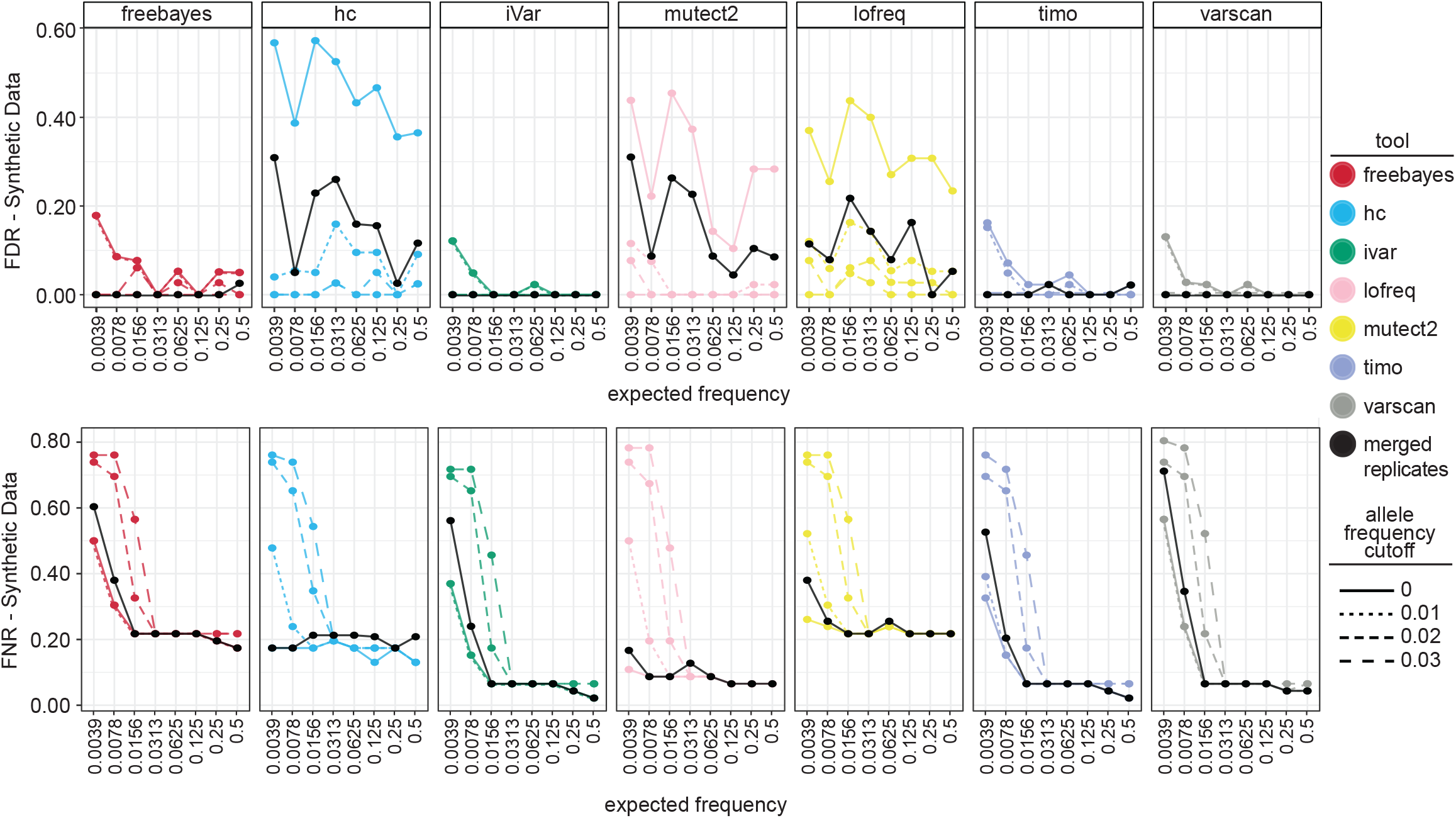
Effect of frequency cutoffs and sequencing replicates on false discovery rate and false negative rate in synthetic and simulated data. **(A)** False discovery rate (FDR) of synthetic data using either single replicate data (colored points and lines) with applied frequency cutoffs (line type) or merged two replicate data without cutoffs (solid black points and lines). Lines represent mean across viruses. **(B)** False negative rate (FNR) of synthetic data using either single replicate data (colored points and lines) with applied frequency cutoffs (line type) or merged two replicate data without cutoffs (solid black points and lines) as above.

Together, these results indicate that using replicate sequencing with less stringent frequency cutoffs (AF >= 0.01) is the best combination to reduce the FDR while maintaining a low FNR. However, when replicate sequencing is unavailable, high coverage across the genome and strict frequency cutoffs (AF >= 0.03) are necessary.

### Choice of variant caller significantly impacts set and frequency of identified variants in real SARS-CoV-2 data using single replicate data

While simulated and synthetic data allow for testing minority variant callers and cutoffs in a controlled setting, real data will always be more unpredictable. Thus, after using simulated and synthetic data to assess variant caller performance across frequencies and coverages, we tested how the callers performed on SARS-CoV-2 sequence data from diagnostic samples. Based on the simulated and synthetic data testing, we determined that a coverage cutoff of 200X and an allele frequency cutoff of 0.03 in single replicate data minimized false positive calls without sacrificing large amounts of true positive data with most variant calling tools. To test the variant calling tools on high-quality data, we used only samples where at least 80% of the genome had a read depth over 200x coverage cutoff in both sequencing replicates (**Fig. S7A**). We used each variant calling tool to identify minority variants in these samples and filtered them using a read depth cutoff of 200 and an allele frequency cutoff of 0.03.

We were interested in how similar the set of variants was that was identified by each caller. As a proof of principle, we filtered the set of variants for those present above an allele frequency of 0.5 and at read depths greater than 5x to identify consensus changes (AF >=50%, or major variants) within the data. As expected, the tools largely agreed on the consensus changes within the data (**Fig. S7B**). There was a small set of major variants that the callers did disagree on, however, most of which were a result of differences in the way some callers identify indels or handle variant at consecutive nucleotide positions. For the purposes of this study, indels were excluded from the analysis. These data indicate that even at high allele frequencies, the variant callers disagree to some extent on the set of variants present in clinical data, an important consideration when choosing how to define consensus sequences from SARS-CoV-2 data.

We then analyzed the intersection of the minority variants (allele frequencies between 0.03 and 0.5) identified by each tool. The total number of variants identified varied greatly between the callers, with Varscan calling the fewest variants by far, followed by timo and Lofreq, in line with the more conservative nature of these callers observed in the previous analyses (**Fig. 4A**). Of note, we found that replicate 2 data had much higher numbers of minority variants, particularly at very low frequencies, regardless of Ct value or date of sequencing. This suggests that freeze thawing samples may impact minor variant numbers (**Fig. S7C**). When comparing the set of minority variants identified by each of the seven tools, there was significant disagreement between the variants. Mutect2 and HaplotypeCaller identified many variants that other callers did not, particularly in replicate 1, and missed several variants identified by the other callers. This was similar to the performance of these callers on the synthetic data sets (**Fig. S7D**). Given the high number of false positives identified by HaplotypeCaller, Mutect2 and Lofreq in the simulated and synthetic datasets, we focused on the intersection of minority SNVs found in just the other four variant callers: Freebayes, iVar, Varscan, and timo. Of all the minority variants found in the data, 104 from replicate 1, and 142 from replicate 2 were identified by all four of the variant callers (**Fig. 4B**). Overall, choice of variant caller appears to have a significant impact on the set of minority variants identified in SARS-CoV-2 data from clinical specimens.

**Figure 4.**
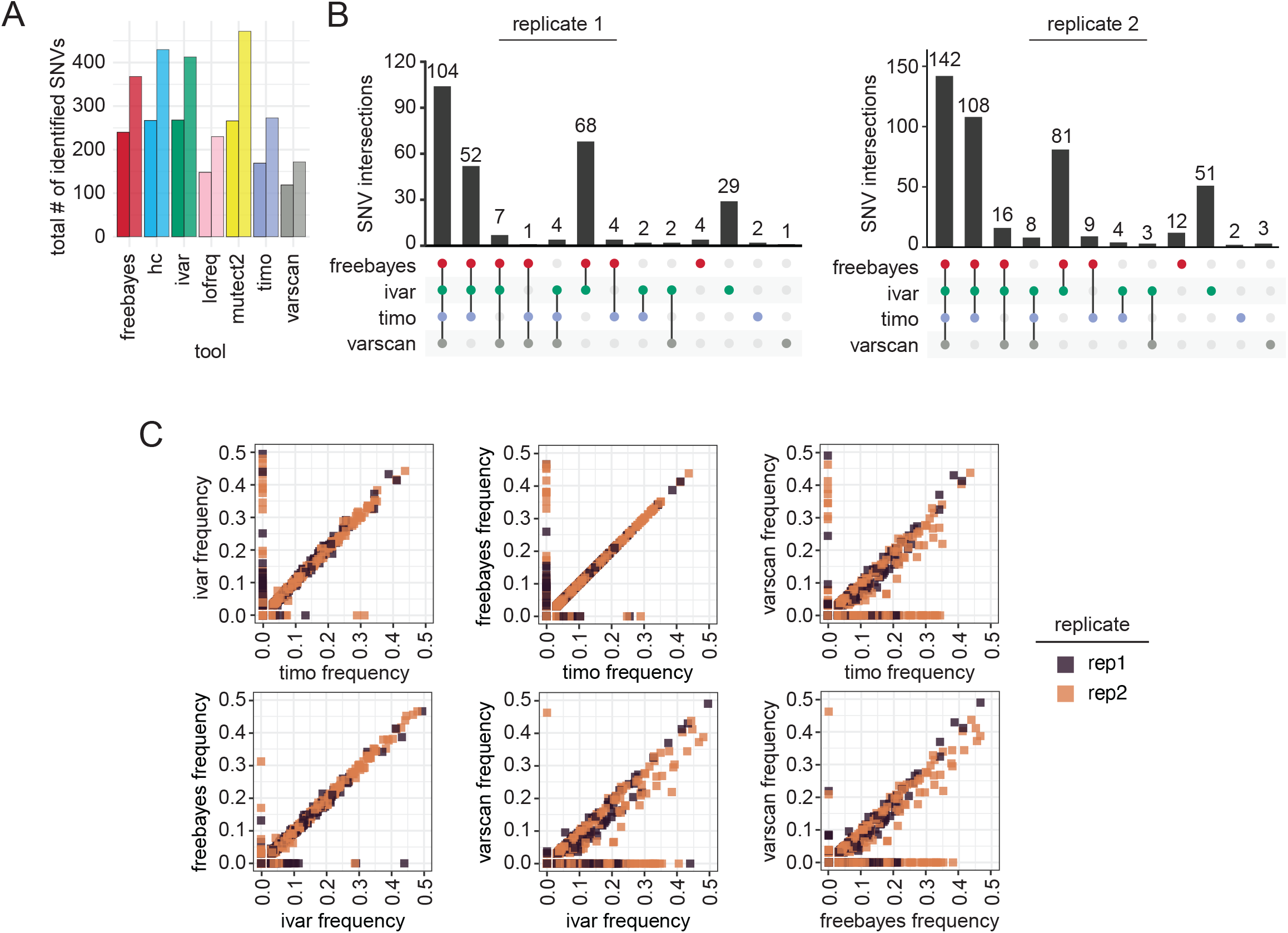
Effect of variant caller on identification and allele frequency estimation of SNVs in SARS-CoV-2 data from clinical samples. **(A)** Bar plot showing raw number of variants identified by each variant caller in replicate 1 (left bar) or replicate 2 (right bar). **(B)** Upset plot showing agreement of minority variants between variant callers in each replicate using an allele frequency cutoff of 0.03 and coverage cutoff of 200X. Vertical bars indicate the size of the shared set of variants while dots and connecting lines show which callers share a given set of identified variants. **(C)** Scatter plot showing the output frequency of minority variants identified by two different variant callers. Color represents replicate. Variants with frequency of 0 were not identified by that variant caller.

Many studies of minority variants investigate frequency of minority variants to calculate selection, bottleneck size, and potential for transmission (24, 41). We were interested in how well the variant callers agreed on the frequency at which variants were identified. We plotted the frequency of a variant in one caller against the frequency in each other caller and found that most of the minority variant callers were strikingly similar in their frequency calls of shared variants. Timo, Freebayes and iVar all showed almost complete agreement on frequency of the variants (**Fig. 4C**). Varscan showed more variation in frequency, generally calling variants at a lower frequency than the other three tools (**Fig. 4C**). Of interest, variants called by one caller but not another spanned a frequency range of 0.03 all the way to 0.5, indicating that even high frequency minority variants were often not agreed upon by variant callers. These data show that choice of variant caller not only affects the set of the minority variants that are identified in a data set, but also the frequency of those variants.

### Most minority variants in data from SARS-CoV-2 clinical specimens are not reproducible across sequencing replicates

In our sequencing data, the number of variants identified in each replicate by each tool was markedly different, suggesting that many of the identified minor variants may not be true variants introduced through viral replication, but instead technical artifacts (**Fig. 4A, S7C-D**). As was shown with our simulated and synthetic data, errors introduced through PCR, library preparation, and sequencing are mostly random, and therefore less likely to reappear and be identified across multiple sequencing replicates, particularly when using Freebayes, iVar, timo, or Varscan (**Fig. 3**). To find high confidence minority variants, we looked at the intersection of variants between the two replicates using each caller, using a lower allele frequency threshold, 0.01, as established in synthetic data for merged replicates (**Fig. 3**). iVar and Freebayes called the highest number of reproducible variants, while timo called the fewest number of reproducible variants, however the percentage of reproducible variants compared to variants identified in single replicates was the highest (**Fig. 5A**). Timo is one of the most conservative callers that was tested; these data indicate that being conservative may lead to increased reproducibility and therefore increased confidence when used on single replicate data. It is however important to note that the relatively low percentages of reproducible variants are likely skewed by the high numbers of low frequency variants found in replicate 2 (**Fig. 5A, S7C**). When we looked at the intersection of only the variants found by each tool in both replicates, only 80 SNVs were found by all callers across replicates, suggesting again that variant callers do not agree on the set of minority variants present (**Fig. 5B**). Together, these data suggest that most minority variants are not reproducible across replicates and support the idea that more than any other criteria, sequencing replicate has the highest impact on the set of minority variants identified (**Fig. 5B, S7C-D**).

**Figure 5.**
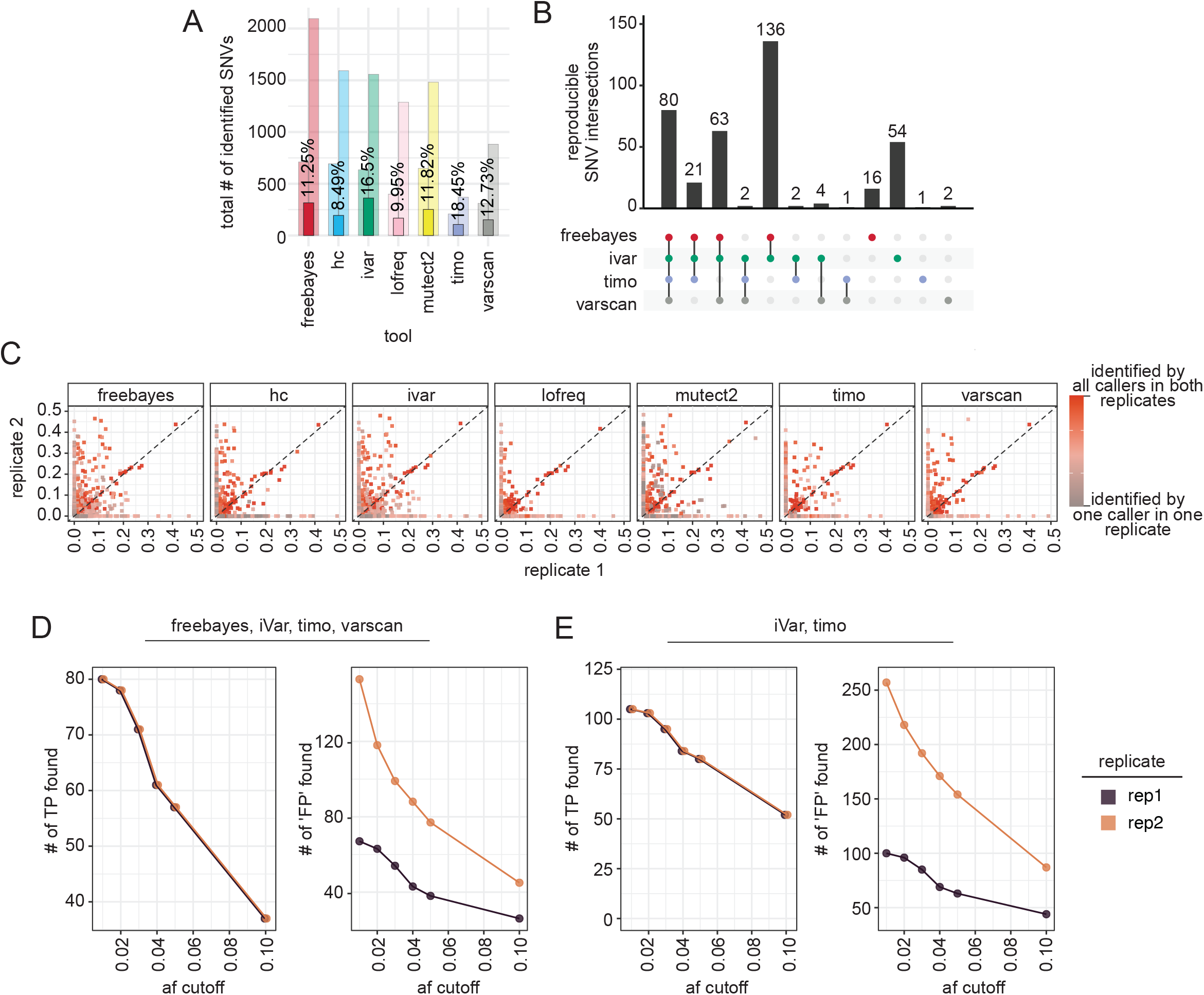
Reproducibility of minority variants across sequencing replicates. **(A)** Bar plot showing number of reproducible variants across sequencing replicates by each variant caller. Percentages shown are the percentage of total individual variants that were reproducible. Background bars indicate the total number of variants found by each tool in each replicate (r1-left, r2-right). **(B)** UpsetR plot showing overlap of reproducible variants across Freebayes, iVar, timo, and Varscan, using a frequency cutoff of 0.01 and coverage cutoff of 200x. Vertical bars indicate the size of the shared set of variants while dots and connecting lines show which callers share a given set of reproducible variants. **(C)** Scatter plot showing frequency of variants across sequencing replicates with frequency in replicate 1 on the x-axis and frequency in replicate 2 on the y-axis. Color represents reproducibility of each variant across variant callers and replicates. **(D**,**E)** Line graph showing the number of ‘true positive’ (D) and ‘false positive’ (E) in single replicate data across allele frequency cutoffs compared to merged replicate data. A TP variant is defined as a SNV found by the selected callers in both replicates (80 variants shown in **B**) and a FP is defined as any other variant found in an individual replicate by the selected callers. Color represents sequencing replicate.

### Variants identified by all variant callers show the most reproducible frequencies

Using synthetic data, we showed that in a controlled setting, SNVs that were found in both sequencing replicates generally showed reproducible frequencies (**Fig. 2C**). Given that frequency is an important metric in most analyses performed using minority variant data, we wanted to test if this held true in clinical samples. Plotting the frequency in replicate 1 against the frequency in replicate 2 revealed that while some variants showed consistent frequencies, some differed drastically—identified at 5-10% in one replicate and as high as 45-50% in the other replicate (**Fig. 5C**). These data were striking as they reveal that averaging frequency across replicates, or performing only one sequencing replicate, could drastically alter downstream analyses performed using these numbers. Interestingly, when we looked at the variants that were reproducible across replicates and found by most, or all the variant callers, frequency tended to be much more consistent than those identified only in a single replicate, or by a single caller (**Fig. 5C**, dark red points). Together, these data suggest that confidence in each variant and its frequency is increased with replicate sequencing and identification by many variant callers.

Since replicate sequencing data is not always available, we investigated what frequency cutoff could be applied such that single replicate data closely resembled the merged replicate data. To do this, we looked at the intersection of SNVs called in both replicates by Freebayes, iVar, timo, and Varscan (80 variants shown in **Fig. 5B**) and compared those to the intersection of SNVs called by the same four callers in each individual replicate (**Fig. 4A**). We then applied allele frequency cutoffs between 0.01 and 0.1 to determine the best cutoff for use on single replicate data. Here, we identify a true positive as a variant present in the reproducible set, and a false positive as any other variant found in a single replicate. As was noted previously, we find that replicate 2 data shows an increased number of SNVs, perhaps due to freeze/thawing of samples between preparations (**Fig. S7C, 5D**). As such, replicate 1 is likely more representative of what single replicate data may typically look like. At an allele frequency cutoff of 0.01, all true positives were found, but the number of false positives was very high, while a frequency cutoff of 0.05 or removed an outsized number of true positives from the dataset (**Fig. 5D**). Based on these data, we suggest an allele frequency cutoff of 0.03 when only single replicate data is available, a cutoff that was also confirmed in the simulated and synthetic data sets (**Fig. 3**). We further suggest using the intersection of multiple variant callers to increase confidence in the data, especially when estimating SNV frequency (**Fig. 5B**). Using all variant callers for analysis would likely be tedious and unrealistic, thus we looked at the intersection of just two callers, iVar and timo, and we found a similar trade-off in true positive and false positive data when using a single replicate and a cutoff of 0.03 (**Fig. 5E**). Based on these data, it is clear that there are many considerations necessary when performing minority variant analyses, and parameters and cutoffs should thus be chosen carefully and thoughtfully depending on the data available. In general, using replicate data and multiple callers provides the highest confidence set of SNVs and the most accurate frequencies.

## DISCUSSION

It has long been understood that intrahost viral populations are heterogeneous in nature, however capturing and measuring this viral diversity is complicated due to errors introduced during preparation and sequencing. We set out to identify the optimal tools, parameters, and filtering methods necessary for accurate variant identification. To accomplish this goal, we used a combination of simulated and synthetic influenza and SARS-CoV-2 sequence data to test the technical and experimental challenges and limitations of minority variant analyses. We found that sequencing depth and choice of variant caller has a significant impact on sensitivity of minor variant calls, and most false positive SNVs were detected at either low allele frequency, or low read depth. Additionally, our results show that replicate sequencing allows for the use of lower frequency thresholds, and this combination provides the best results, keeping the false discovery rate low, without sacrificing true positive data. We tested our optimized frequency and coverage cutoffs using SARS-CoV-2 sequence data from clinical infections. Most variant callers did not agree on the set of minor variants in the virus sequence data from clinical samples, and most minority variants were not reproducible across replicates. Ultimately, we determined that using a combination of sequencing replicates and multiple variant callers, along with moderate allele frequency and coverage cutoffs, can increase confidence in SNV calls. Our results outline the main considerations for minority variant analyses and highlight the intricacies and difficulties of studies of this nature. Further, we provide a framework for designing minority variant analyses, which will ultimately lead to more accurate conclusions surrounding viral transmission and evolution.

Using a standardized set of parameters, most callers performed relatively similarly on high coverage simulated data, having both high precision and high recall. The main differences in caller performance were seen in lower coverage data or at low frequencies. As many minority variants are found at low frequencies, understanding how tools perform under these conditions is more relevant to analyses of real sequencing data. Timo had the lowest recall at lower coverages and simulated frequencies due to its rigid requirements for SNVs to be above the 0.01 threshold parameter, while many other callers found SNVs at or below this frequency, regardless of setting a 0.01 AF cutoff. Timo, iVar, and Varscan all have the functionality to drop the input frequency parameter down to 0.001. Decreasing this parameter did not change the accuracy of iVar and Varscan but did increase the recall of timo. These data highlight the importance of optimizing bioinformatic tools to one’s own data.

As previously observed by our group and others, the best method for filtering out errors generated during sample processing is to sequence each sample twice and only keep the SNVs found in both replicates. Sequencing replicates removed nearly all false positive calls in simulated data and significantly reduced the number of false positive SNVs in the synthetic datasets. However, for the synthetic datasets, the number of false positive SNVs was highly dependent on the variant caller used. HaplotypeCaller, Lofreq, and Mutect2 were all made and optimized for identifying variants in cancer cell datasets and had significantly higher false discovery rates than tools designed for viral use, particularly at low allele frequencies. Adjusting the filtering or input parameters on these callers may better optimize them for their use on viral data. For example, HaplotypeCaller suggests additional filtering of output data, however, when performed on this dataset, SNV detection was significantly reduced. Without this additional filtering, most variants are identified (high recall) but high numbers of false positives are included, suggesting additional optimization could improve performance.

The use of simulated and synthetic data allows tools to be tested against a true set of minority variants. This is invaluable given that it is not possible to know which SNVs are real, and which are not, in clinical sequencing data. However, each of these controlled settings comes with both benefits and limitations. Simulated data allows for testing of various coverages using random downsampling, showing how low coverage data impacts caller performance and allele frequency estimation. Since high coverage is often the goal in sequencing, and coverage is unpredictable with real samples (42), this is a variable that can only be truly tested through simulation.

Conversely, while error models in simulated data can imitate PCR and sequencing errors, there are limitations to how ‘real’ these errors look to variant callers. Synthetic data allows for real error to be introduced during sample preparation and ultimately be easily separated from real variation during analysis. This gives a more accurate depiction of false positives that would exist in clinical data and variant caller performance on synthetic data may be more representative of true performance. However, both methods are limited in their ability to mimic sequence data from clinical samples where the RNA is not as homogeneous and intact as with synthetic RNA, and likely contains human contamination from sample collection and potential degradation from freezing, thawing, and handling (43). This is evident in the disagreement between variant callers and replicates, both on the set of minority variants and on their frequencies, in real data. Combined, the simulated, synthetic, and clinical data sets show that there will always be a trade-off between inclusion of the maximum number of true variants, and inclusion of false positive data.

Our study provides an extensive framework for studying minority variants in sequence data from clinical samples, outlining major considerations around choice of variant caller, application of frequency and coverage thresholds, and use of replicate sequencing. Further, we have established a pipeline that can be used for further testing and optimization of parameters, or for other viruses. This work will inform and improve future studies of intrahost variation and estimates surrounding viral diversity and viral evolution.

## SUPPLEMENTAL TABLES

**Supplemental Table 1**. List of variant callers and parameters used for each. **Supplemental Table 2**. Metadata associated with diagnostic samples processed for whole genome sequencing of SARS-CoV-2.

## SUPPLEMENTAL FIGURES

**Supplemental Figure 1. Experimental setup of simulated and synthetic data generation. (A)** Schematic of the SNV simulation pipeline. **(B)** Nucleotide positions of SNVs in simulated data across the genomes of A/H1N1 (n=121), A/H3N2 (n=110), B/Victoria (n=118), and SARS-CoV-2 (n=144). Gene segments for influenza A and B strains are ordered largest (PB2) to smallest (NS), left to right. **(C)** Schematic of allele frequency and viral loads used for mixing WT and variant RNA for library preparation and sequencing. **(D)** Location of synthetic SNVs across the three influenza gene segments. Gray lines represent the designed SNVs (PB2 (n=18), HA (n=14), NA (n=14)) and red lines represent the pre-mRT-PCR errors, likely generated during template preparation or *in vitro*-transcription.

**Supplemental Figure 2. Variant caller performance using default, standard, and custom parameters. (A)** F1 statistic for each variant caller across a range of downsampling fractions and allele frequencies. Values shown are mean and standard deviation of four viruses (A/H1N1, A/H3N2, B/Victoria, SARS-CoV-2) using default (top), standard (middle) or custom (bottom) input parameters (**Table S1**). Color represents the variant caller used. Dark timo points and lines indicate timo output with the removal of the binomial check option.

**Supplemental Figure 3. Coverage and Expected vs. Observed frequency of variants in synthetic data samples (A)** Coverage plots showing log10 read depth for each synthetic influenza control across the PB2, HA, and NA gene segments. Color represents individual samples. **(B)** Expected (x-axis) versus observed frequency (y-axis) for synthetic influenza SNVs (replicate one only). Color represents the copy number. **(C)** Expected (x-axis) versus observed frequency (y-axis) for synthetic SARS-CoV-2 SNVs. Dashed black lines represent y=x.

**Supplemental Figure 4. Effect of frequency cutoffs and sequencing replicates on false discovery rate and false negative rate in simulated data. (A)** False discovery rate (FDR) of simulated data using either single replicate data (colored points and lines) with applied frequency cutoffs (line type) or merged two replicate data without cutoffs (solid black points and lines). Background points represent values for individual viruses. Lines represent mean across viruses. **(B)** False negative rate (FNR) of simulated data using either single replicate data (colored points and lines) with applied frequency cutoffs (line type) or merged two replicate data without cutoffs (solid black points and lines) as above.

**Supplemental Figure 5. Effect of coverage cutoffs and sequencing replicates on false discovery rate and false negative rate in synthetic and simulated data. (A, B)** False discovery rate (FDR) (**A**) of false negative rate (FNR) (**B**) of synthetic data using either single replicate data (colored points and lines) with applied coverage cutoffs (line type) or merged two replicate data without cutoffs (solid black points and lines). Background points show individual values for each of the four viruses. Lines represent mean across viruses. (**C, D**) False discovery rate (FDR) (**C**) of false negative rate (FNR) (**D**) of synthetic data using either single replicate data (colored points and lines) with applied coverage cutoffs (line type) or merged two replicate data without cutoffs (solid black points and lines) as above.

**Supplemental Figure 6. Effect of sequencing replicates on the accuracy of allele frequency estimation. (A)** Scatter plot showing the frequency of variants in synthetic influenza data across sequencing replicates with frequency in replicate 1 on the x-axis and frequency in replicate 2 on the y-axis. Color represents the SNV type. **(B, C)** Percent error 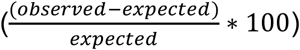 of single replicate (greyed boxes) or merged replicate (open boxes) SNVs in simulated data across downsampling fractions (**B**) or synthetic data across expected frequencies and viral loads (**C**). Color represents the variant caller used.

**Supplemental Figure 7. Quantification of majority and minority variants identified in data from SARS-CoV-2 clinical specimens (A)** Scatter plot showing Ct value against percentage of genome with coverage over 200x after filtering for only samples with 80% of the genome over the 200x cutoff. **(B)** Upset plot showing agreement of consensus changes between variant callers in each replicate using an allele frequency cutoff of 0.5 and coverage cutoff of 5X. Vertical bars indicate the size of the shared set of changes while dots and connecting lines show which callers share a given set of identified changes. **(C)** Box and whisker plot showing number of minor variants with indicated allele frequency cutoffs found in replicate 1 and replicate 2 sequencing data. Points represent individual samples. Boxes and whiskers show min, first quartile, median, third quartile and max for each replicate. **(D)** Upset plot showing agreement of minority variants between all variant callers in each replicate using an allele frequency cutoff of 0.03 and coverage cutoff of 200x. Vertical bars indicate the size of the shared set of variants while dots and connecting lines show which callers share a given set of identified variants.

## REFERENCES

1. Arnold JJ, Cameron CE. 2004. Poliovirus RNA-dependent RNA polymerase (3Dpol): presteady-state kinetic analysis of ribonucleotide incorporation in the presence of Mg2+. Biochemistry 43:5126–37.

2. Sanjuan R. 2012. From molecular genetics to phylodynamics: evolutionary relevance of mutation rates across viruses. PLoS Pathog 8:e1002685.

3. Duffy S, Shackelton LA, Holmes EC. 2008. Rates of evolutionary change in viruses: patterns and determinants. Nat Rev Genet 9:267–76.

4. Peck KM, Lauring AS. 2018. Complexities of Viral Mutation Rates. J Virol 92.

5. Domingo E. 2002. Quasispecies Theory in Virology. J Virol 76:463–465.

6. Sanjuan R, Domingo-Calap P. 2016. Mechanisms of viral mutation. Cell Mol Life Sci 73:4433–4448.

7. Lynch M, Ackerman MS, Gout JF, Long H, Sung W, Thomas WK, Foster PL. 2016. Genetic drift, selection and the evolution of the mutation rate. Nat Rev Genet 17:704–714.

8. Gorbalenya AE, Enjuanes L, Ziebuhr J, Snijder EJ. 2006. Nidovirales: evolving the largest RNA virus genome. Virus Res 117:17–37.

9. Smith EC, Sexton NR, Denison MR. 2014. Thinking Outside the Triangle: Replication Fidelity of the Largest RNA Viruses. Annu Rev Virol 1:111–32.

10. Sanjuan R, Moya A, Elena SF. 2004. The distribution of fitness effects caused by singlenucleotide substitutions in an RNA virus. Proc Natl Acad Sci U S A 101:8396–401.

11. Visher E, Whitefield SE, McCrone JT, Fitzsimmons W, Lauring AS. 2016. The Mutational Robustness of Influenza A Virus. PLoS Pathog 12:e1005856.

12. Peck KM, Chan CH, Tanaka MM. 2015. Connecting within-host dynamics to the rate of viral molecular evolution. Virus Evol 1:vev013.

13. Wang Y, Wang D, Zhang L, Sun W, Zhang Z, Chen W, Zhu A, Huang Y, Xiao F, Yao J, Gan M, Li F, Luo L, Huang X, Zhang Y, Wong SS, Cheng X, Ji J, Ou Z, Xiao M, Li M, Li J, Ren P, Deng Z, Zhong H, Xu X, Song T, Mok CKP, Peiris M, Zhong N, Zhao J, Li Y, Li J, Zhao J. 2021. Intra-host variation and evolutionary dynamics of SARS-CoV-2 populations in COVID-19 patients. Genome Med 13:30.

14. Martin MAK, Katia. 2021. Reanalysis of deep-sequencing data from Austria points towards a small SARS-COV-2 transmission bottleneck on the order of one to three virions. bioRxiv doi:https://doi.org/10.1101/2021.02.22.432096.

15. McCrone JT, Lauring AS. 2018. Genetic bottlenecks in intraspecies virus transmission. Curr Opin Virol 28:20–25.

16. Wang D, Wang Y, Sun W, Zhang L, Ji J, Zhang Z, Cheng X, Li Y, Xiao F, Zhu A, Zhong B, Ruan S, Li J, Ren P, Ou Z, Xiao M, Li M, Deng Z, Zhong H, Li F, Wang WJ, Zhang Y, Chen W, Zhu S, Xu X, Jin X, Zhao J, Zhong N, Zhang W, Zhao J, Li J, Xu Y. 2021. Population Bottlenecks and Intra-host Evolution During Human-to-Human Transmission of SARS-CoV-2. Front Med (Lausanne) 8:585358.

17. Lythgoe KA, Hall M, Ferretti L, de Cesare M, MacIntyre-Cockett G, Trebes A, Andersson M, Otecko N, Wise EL, Moore N, Lynch J, Kidd S, Cortes N, Mori M, Williams R, Vernet G, Justice A, Green A, Nicholls SM, Ansari MA, Abeler-Dorner L, Moore CE, Peto TEA, Eyre DW, Shaw R, Simmonds P, Buck D, Todd JA, Oxford Virus Sequencing Analysis G, Connor TR, Ashraf S, da Silva Filipe A, Shepherd J, Thomson EC, Consortium C-GU, Bonsall D, Fraser C, Golubchik T. 2021. SARS-CoV-2 within-host diversity and transmission. Science doi:10.1126/science.abg0821.

18. Pybus OG, Rambaut A. 2009. Evolutionary analysis of the dynamics of viral infectious disease. Nat Rev Genet 10:540–50.

19. Rockett R, Basile K, Maddocks S, Fong W, Agius JE, Johnson-Mackinnon J, Arnott A, Chandra S, Gall M, Draper J, Martinez E, Sim EM, Lee C, Ngo C, Ramsperger M, Ginn AN, Wang Q, Fennell M, Ko D, Lim HL, Gilroy N, O’Sullivan MVN, Chen SC, Kok J, Dwyer DE, Sintchenko V. 2022. Resistance Mutations in SARS-CoV-2 Delta Variant after Sotrovimab Use. N Engl J Med 386:1477–1479.

20. Martinez-Gonzalez B, Soria ME, Vazquez-Sirvent L, Ferrer-Orta C, Lobo-Vega R, Minguez P, de la Fuente L, Llorens C, Soriano B, Ramos-Ruiz R, Corton M, Lopez-Rodriguez R, Garcia-Crespo C, Somovilla P, Duran-Pastor A, Gallego I, de Avila AI, Delgado S, Moran F, Lopez-Galindez C, Gomez J, Enjuanes L, Salar-Vidal L, Esteban-Munoz M, Esteban J, Fernandez-Roblas R, Gadea I, Ayuso C, Ruiz-Hornillos J, Verdaguer N, Domingo E, Perales C. 2022. SARS-CoV-2 Mutant Spectra at Different Depth Levels Reveal an Overwhelming Abundance of Low Frequency Mutations. Pathogens 11.

21. Dinis JM, Florek KR, Fatola OO, Moncla LH, Mutschler JP, Charlier OK, Meece JK, Belongia EA, Friedrich TC. 2016. Deep Sequencing Reveals Potential Antigenic Variants at Low Frequencies in Influenza A Virus-Infected Humans. J Virol 90:3355–65.

22. Xue KS, Bloom JD. 2020. Linking influenza virus evolution within and between human hosts. Virus Evol 6:veaa010.

23. Valesano AL, Taniuchi M, Fitzsimmons WJ, Islam MO, Ahmed T, Zaman K, Haque R, Wong W, Famulare M, Lauring AS. 2021. The Early Evolution of Oral Poliovirus Vaccine Is Shaped by Strong Positive Selection and Tight Transmission Bottlenecks. Cell Host Microbe 29:32–43 e4.

24. Lauring AS. 2020. Within-Host Viral Diversity: A Window into Viral Evolution. Annu Rev Virol 7:63–81.

25. McCrone JT, Lauring AS. 2016. Measurements of Intrahost Viral Diversity Are Extremely Sensitive to Systematic Errors in Variant Calling. J Virol 90:6884–95.

26. Valesano ALR, Kalee E; Dimcheff, Derek E; Blair, Christopher N; Fitzsimmons, William J; Petrie, Joshua G; Martin, Emily T; Lauring, Adam S. 2021. Temporal dynamics of SARS-CoV-2 mutation accumulation within and across infected hosts. bioRxiv doi:https://doi.org/10.1101/2021.01.19.427330.

27. Grubaugh ND, Gangavarapu K, Quick J, Matteson NL, De Jesus JG, Main BJ, Tan AL, Paul LM, Brackney DE, Grewal S, Gurfield N, Van Rompay KKA, Isern S, Michael SF, Coffey LL, Loman NJ, Andersen KG. 2019. An amplicon-based sequencing framework for accurately measuring intrahost virus diversity using PrimalSeq and iVar. Genome Biol 20:8.

28. Koboldt DC. 2020. Best practices for variant calling in clinical sequencing. Genome Med 12:91.

29. Olson ND, Lund SP, Colman RE, Foster JT, Sahl JW, Schupp JM, Keim P, Morrow JB, Salit ML, Zook JM. 2015. Best practices for evaluating single nucleotide variant calling methods for microbial genomics. Front Genet 6:235.

30. Stead LF, Sutton KM, Taylor GR, Quirke P, Rabbitts P. 2013. Accurately identifying low-allelic fraction variants in single samples with next-generation sequencing: applications in tumor subclone resolution. Hum Mutat 34:1432–8.

31. Auwera Gvd, O’Connor BD. 2020. Genomics in the cloud : using Docker, GATK, and WDL in Terra, First edition. ed. O’Reilly Media, Sebastopol, CA.

32. Cibulskis K, Lawrence MS, Carter SL, Sivachenko A, Jaffe D, Sougnez C, Gabriel S, Meyerson M, Lander ES, Getz G. 2013. Sensitive detection of somatic point mutations in impure and heterogeneous cancer samples. Nat Biotechnol 31:213–9.

33. Garrison EM, Gabor. 2012. Haplotype-based variant detection from short-read sequencing. ArXiv 1207.3907v2.

34. Benjamin D, Sato T, Cibulskis K, Getz G, Stewart C, L. L. 2019. Calling Somatic SNVs and Indels with Mutect2. bioRxiv doi:https://doi.org/10.1101/861054.

35. Koboldt DC, Zhang Q, Larson DE, Shen D, McLellan MD, Lin L, Miller CA, Mardis ER, Ding L, Wilson RK. 2012. VarScan 2: somatic mutation and copy number alteration discovery in cancer by exome sequencing. Genome Res 22:568–76.

36. Wilm A, Aw PP, Bertrand D, Yeo GH, Ong SH, Wong CH, Khor CC, Petric R, Hibberd ML, Nagarajan N. 2012. LoFreq: a sequence-quality aware, ultra-sensitive variant caller for uncovering cell-population heterogeneity from high-throughput sequencing datasets. Nucleic Acids Res 40:11189–201.

37. Wybo WA, Jordan J, Ellenberger B, Marti Mengual U, Nevian T, Senn W. 2021. Datadriven reduction of dendritic morphologies with preserved dendro-somatic responses. Elife 10.

38. Bolger AM, Lohse M, Usadel B. 2014. Trimmomatic: a flexible trimmer for Illumina sequence data. Bioinformatics 30:2114–20.

39. Li H, Durbin R. 2009. Fast and accurate short read alignment with Burrows-Wheeler transform. Bioinformatics 25:1754–60.

40. Van der Auwera GA, Carneiro MO, Hartl C, Poplin R, Del Angel G, Levy-Moonshine A, Jordan T, Shakir K, Roazen D, Thibault J, Banks E, Garimella KV, Altshuler D, Gabriel S, DePristo MA. 2013. From FastQ data to high confidence variant calls: the Genome Analysis Toolkit best practices pipeline. Curr Protoc Bioinformatics 43:11 10 1–11 10 33.

41. Zhao L, Illingworth CJR. 2019. Measurements of intrahost viral diversity require an unbiased diversity metric. Virus Evol 5:vey041.

42. Lambisia AW, Mohammed KS, Makori TO, Ndwiga L, Mburu MW, Morobe JM, Moraa EO, Musyoki J, Murunga N, Mwangi JN, Nokes DJ, Agoti CN, Ochola-Oyier LI, Githinji G. 2022. Optimization of the SARS-CoV-2 ARTIC Network V4 Primers and Whole Genome Sequencing Protocol. Front Med (Lausanne) 9:836728.

43. Li L, Li X, Guo Z, Wang Z, Zhang K, Li C, Wang C, Zhang S. 2020. Influence of Storage Conditions on SARS-CoV-2 Nucleic Acid Detection in Throat Swabs. J Infect Dis 222:203–205.

