## Supplemental Figures for "Optimized Quantification of Intrahost Viral Diversity in SARS-CoV-2 and Influenza Virus Sequence Data"

Supplemental Figure 1

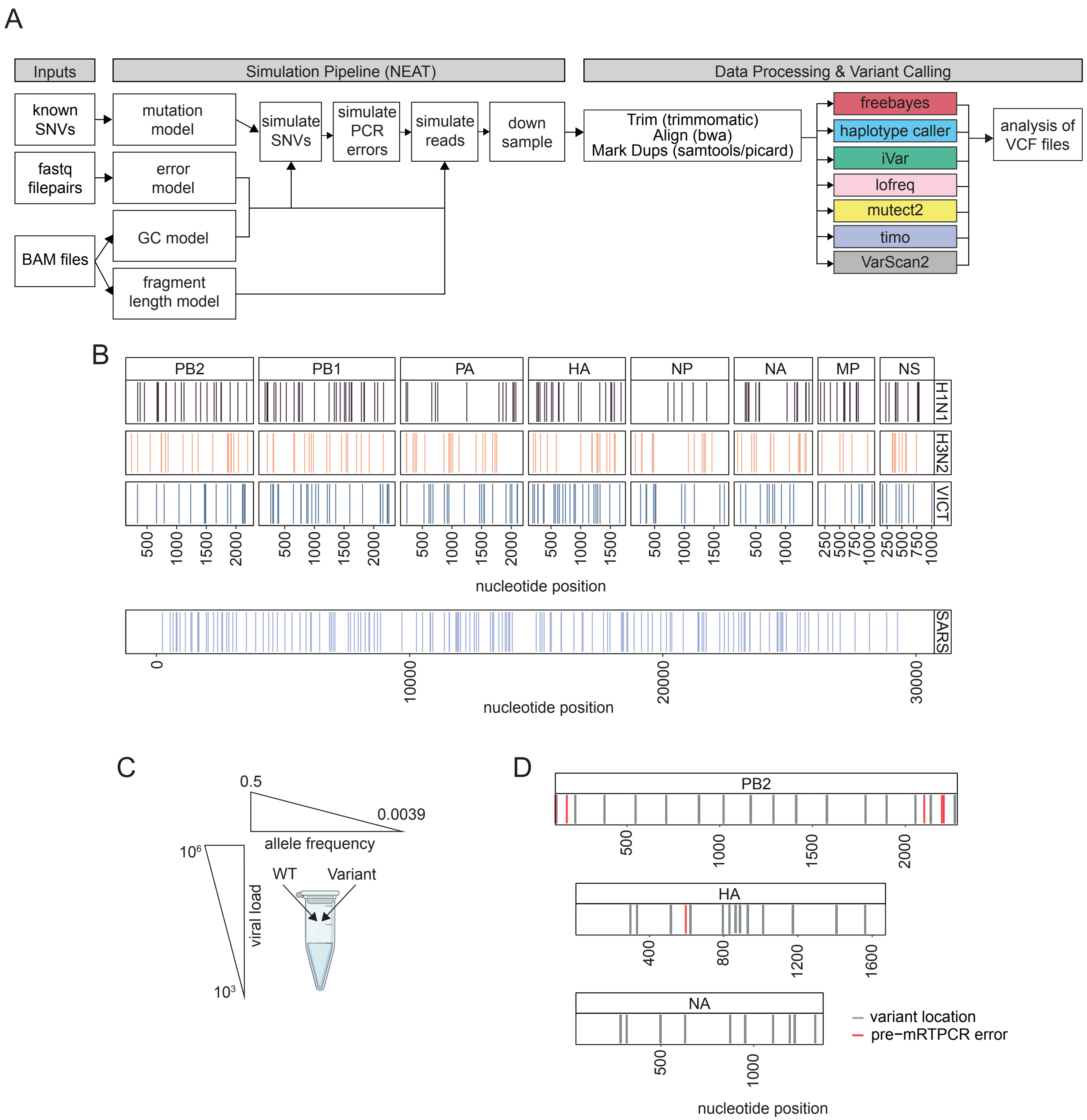

### Supplemental Figure 2

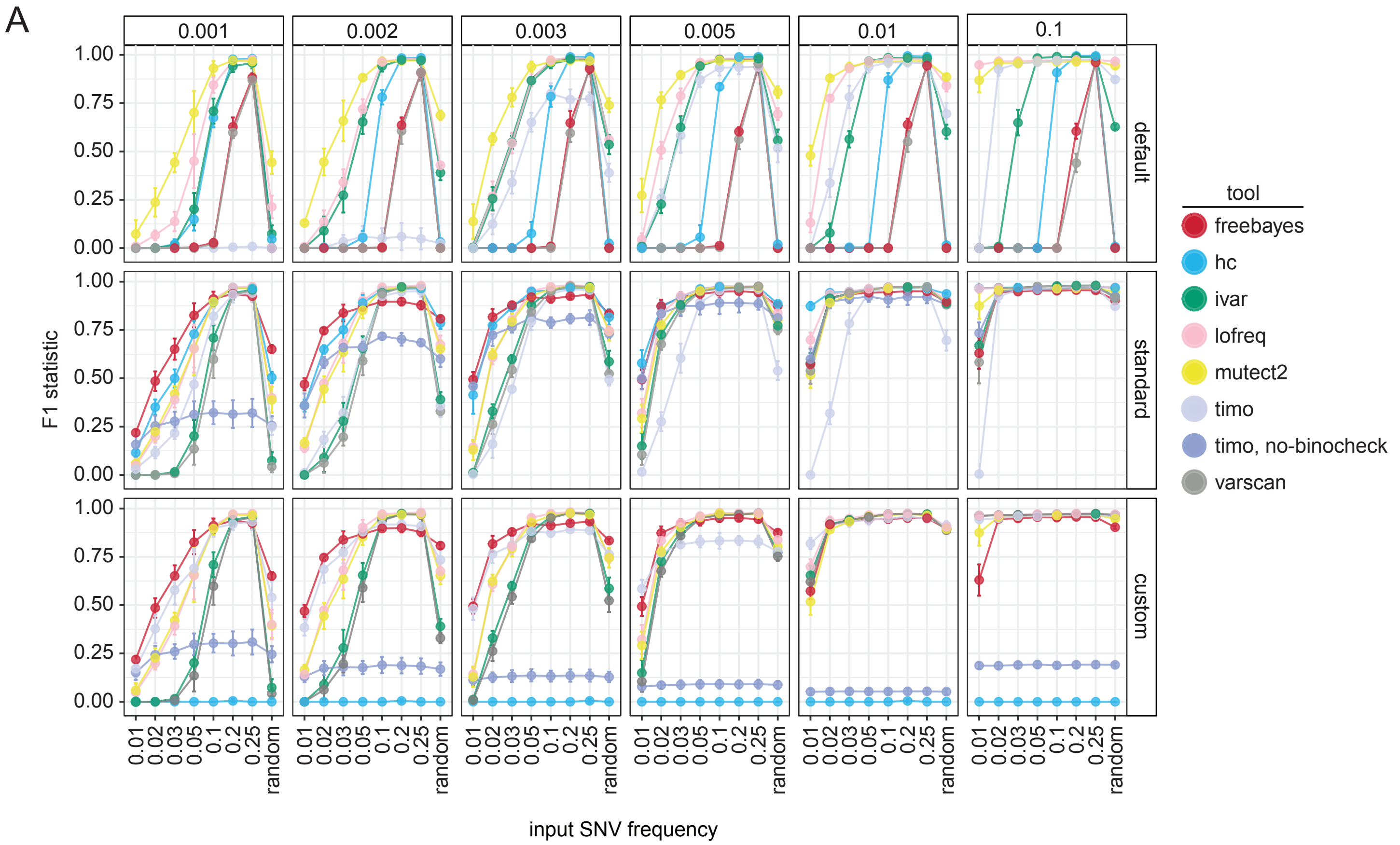

Supplemental Figure 3

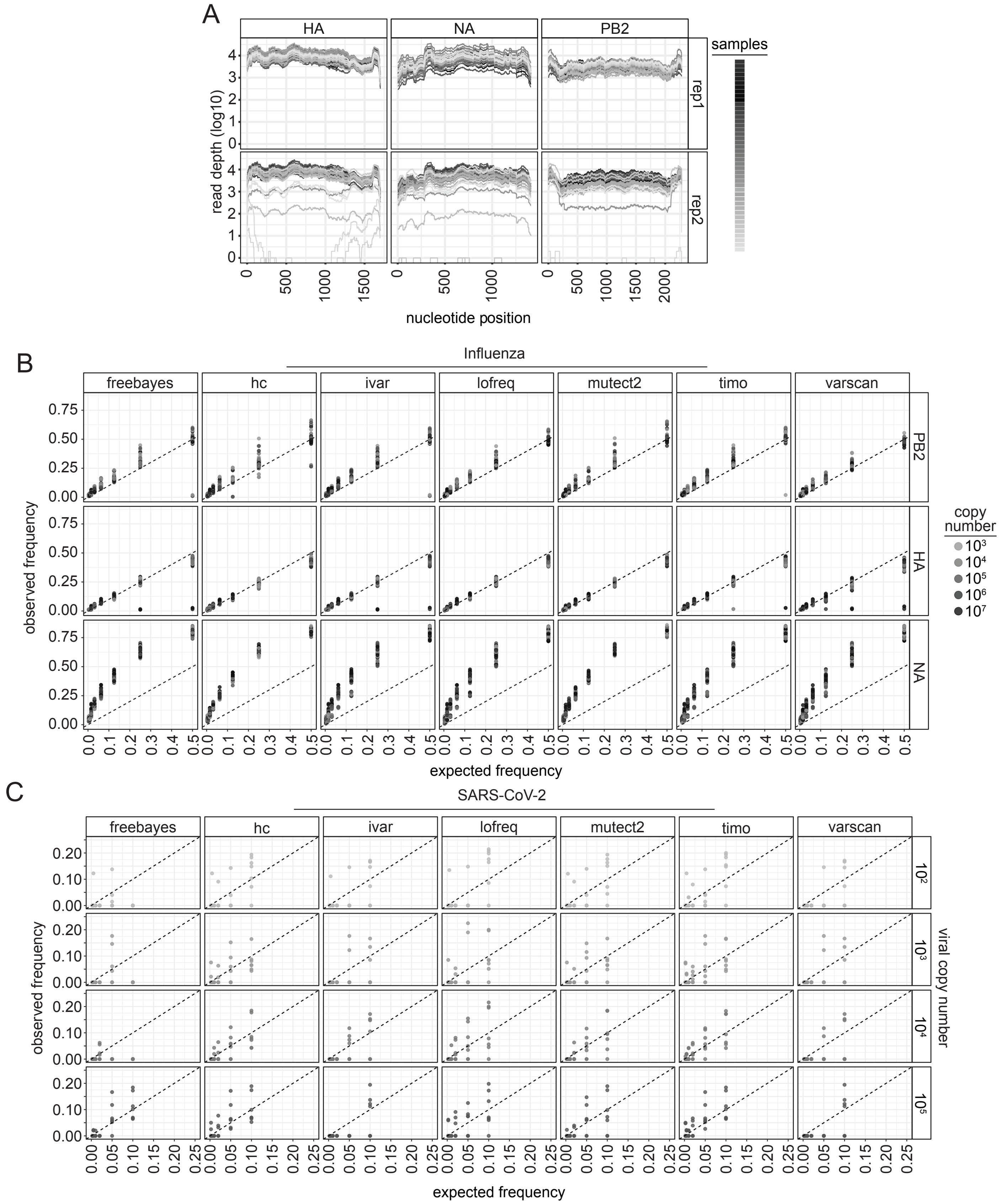

### Supplemental Figure 4

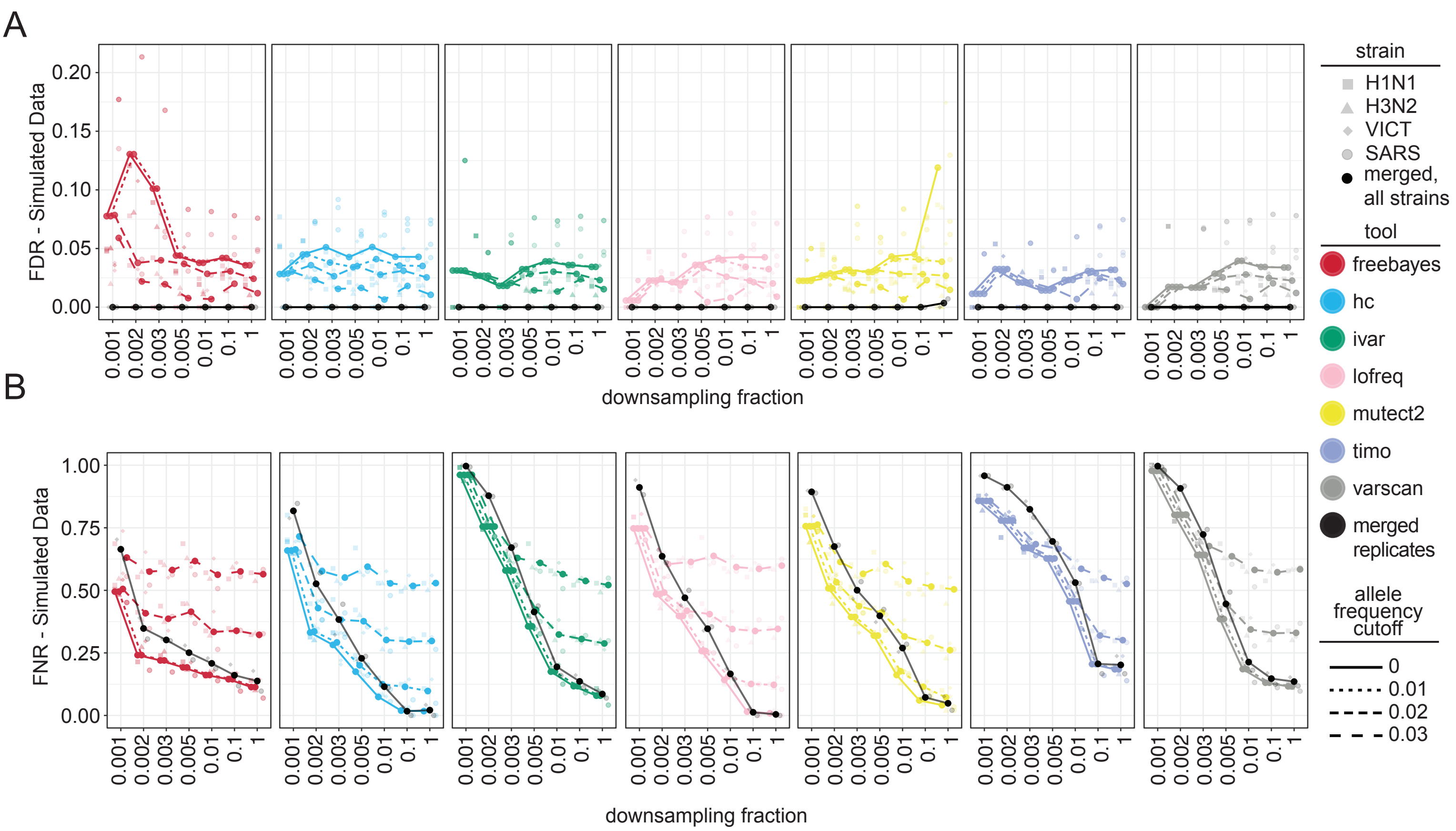

Supplemental Figure 5

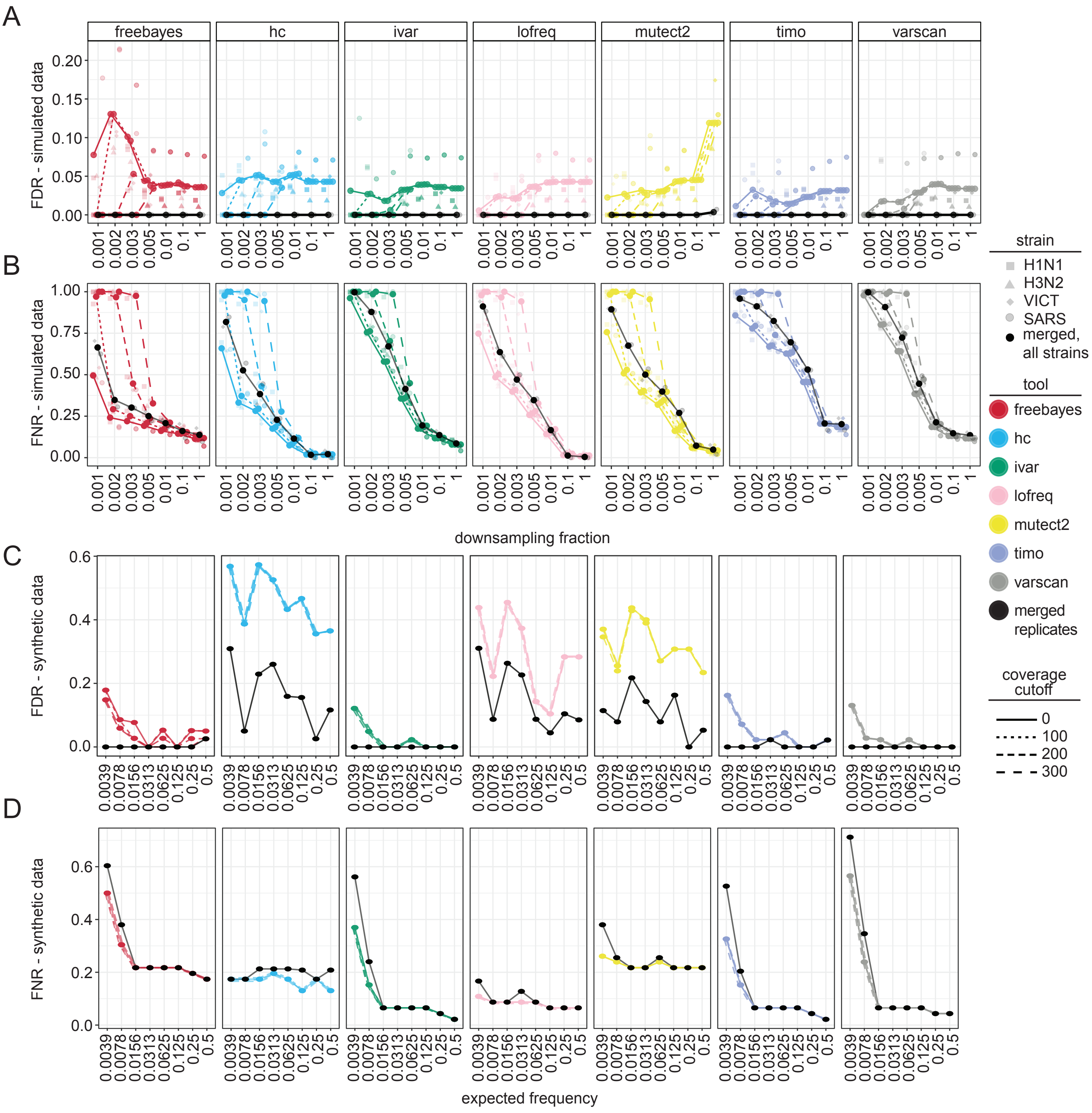

### Supplemental Figure 6

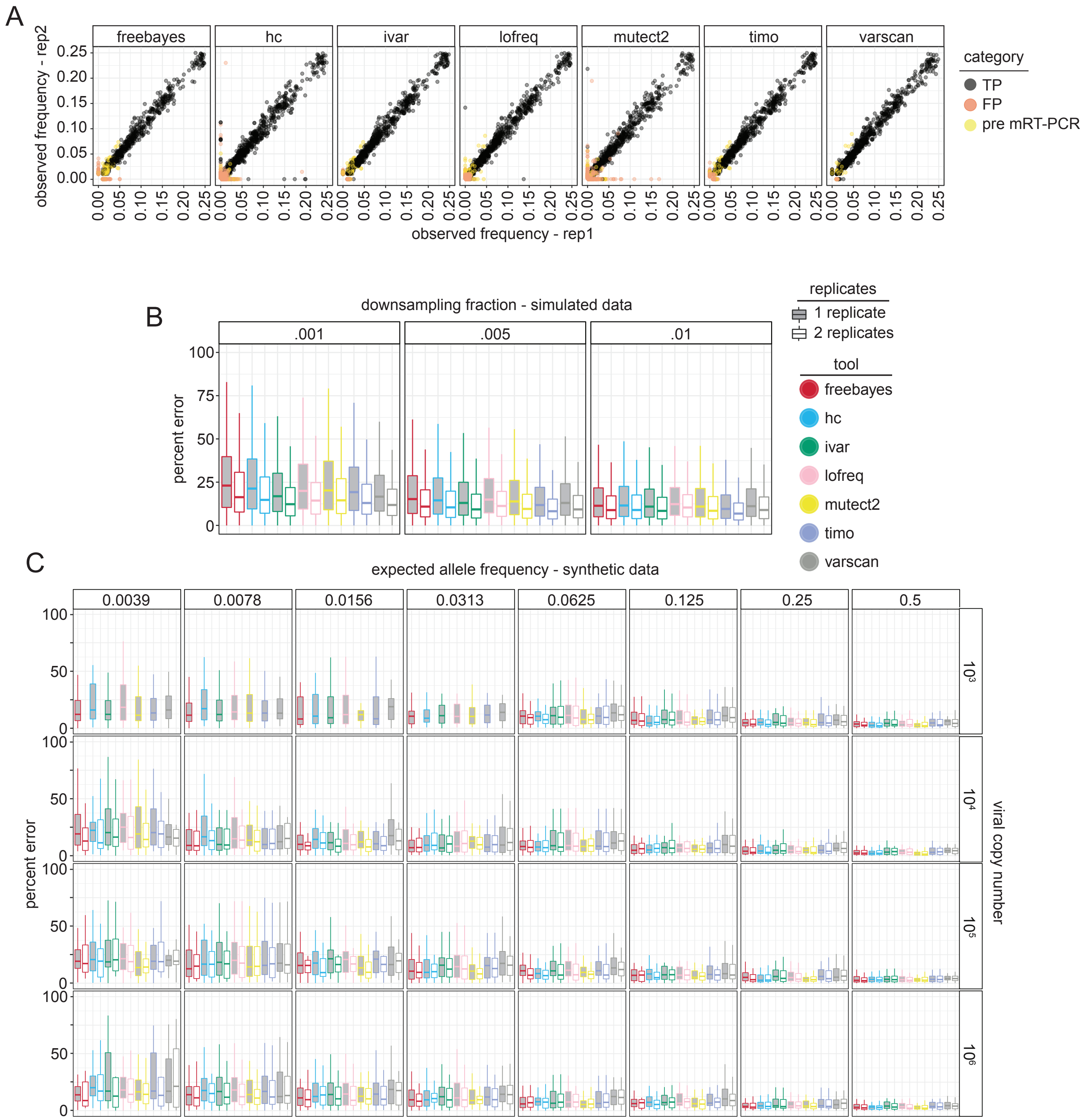

Supplemental Figure 7

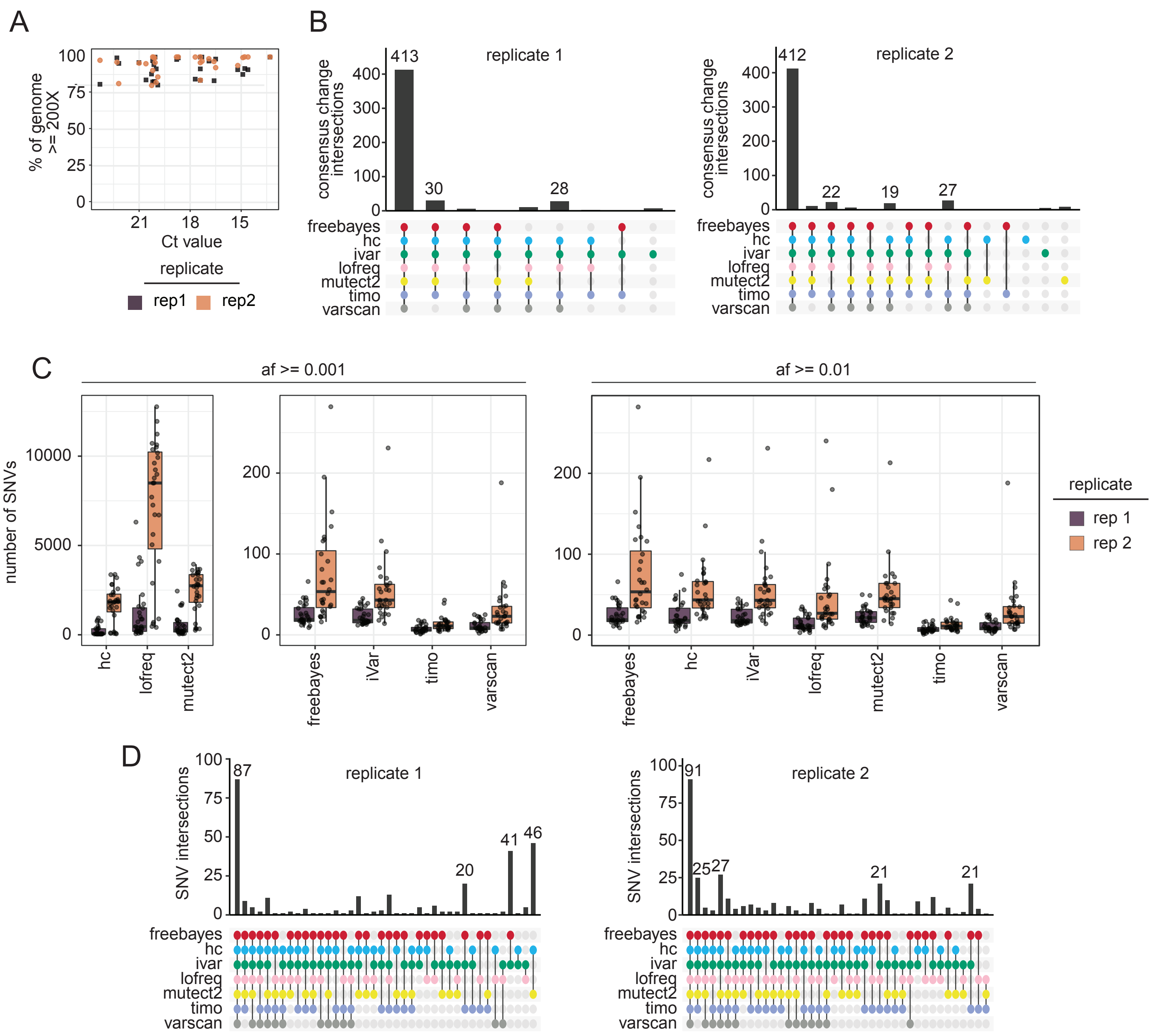
