## Supplemental Methods for "Optimized Quantification of Intrahost Viral Diversity in SARS-CoV-2 and Influenza Virus Sequence Data"

Generation of Simulated Data

Data for the creation of influenza and SARS-CoV-2 mutation models were downloaded from the Influenza Research Database (IRD, www.fludb.org), (downloaded 03/2021) and the China National Center for Bioinformation (CNCB, https://bigd.big.ac.cn/ncov/variation/statistics?lang=en) (downloaded 06/2020) respectively. For influenza, allele counts for all positions in the eight segments of three major influenza virus strains (two subtype of influenza A and one lineage of influenza B) were obtained using the parameters listed in **Table S1**. Custom scripts (available at https://github.com/gencorefacility/MAD2) were used to parse the resulting TSV files to create influenza reference sequences using only the coding sequence. For the MP and NS segments, the sequence between the first stop codon and the last stop codon (i.e. containing both splice forms) was used. Lines were skipped in instances where "N/A" was found as the position within the TSV file with a corresponding "-" as the consensus allele. N's were excluded in determination of the consensus allele. The Wuhan/Hu-1 sequence was used as a SARS-CoV-2 reference (NC_045512.2).

Reads were simulated using NEAT (v2.0) by constructing a mutation, error, fragment length, and GC model for each viral type (1). The mutation model was generated using the VCF and reference FASTA files with the genMutModel.py tool. The sequence error model was created using representative Illumina sequencing data from a paired-end sequencing run of the respective viral subtype and the genSeqErrorModel.py tool. These paired-end sequence reads were also aligned to the custom reference FASTA using BWA mem v0.7.1 (2), and the resulting BAM files were then used to build GC and fragment length models using the computeGC.py and computeFraglen.py tools (NEAT) (1).

These models were then provided to NEAT genReads.py along with the reference fasta files and a mutation rate of 0.009 (0.9%) for influenza and 0.0045 (0.45%) for SARS-CoV-2 to produce a “golden VCF” containing a set number of SNVs in each virus (n=121 (A/H1N1), 110

(A/H3N2), 118 (B/Victoria) and 144 (SARS-CoV-2)). Simulated PCR errors were added using genReads.py (NEAT), using the golden VCF as input and a mutation rate of 0.0005 (0.05%), resulting in 3-8 simulated PCR errors per genome. This process was performed twice, resulting in two replicates with the same set of true SNVs but independently generated PCR mutations.

Several copies of the replicate golden VCFs were made, each with the same variants but with differing allele frequencies (AF): 0.01, 0.02, 0.03, 0.05, 0.1, 0.2, 0.25, 0.5 (one fixed AF per file), and one VCF with Poisson distributed frequencies using a mean of 3, $x \sim \text{Pois}(3)$, and excluding the zero class. Random allele frequencies were programmed to be identical in replicate VCF files by use of a shared seed. For simulated PCR mutations (i.e. those not present in the original golden VCF file), a random allele frequency $x$ was assigned using $x \sim \text{Pois}(2)$, again excluding the zero class. This process resulted in two VCF files containing the common set of golden SNVs at a specified frequency and independent PCR variants at variable frequencies. Each VCF was then provided as input to NEAT genReads.py along with the reference, error model, fragment length model, GC model, and the following parameters: ploidy = 100, read length = 150, coverage = 100,000, and mutation rate = 0.

Each set of simulated paired end fastq libraries was then down-sampled at the following fractions: 0.1 (read depth ~10000x), 0.01 (~1000x), 0.005 (~500x), 0.003 (~300x), 0.002 ~(200x), 0.001 (~100x), 0.0001 (~10x), and 0.00001 (~1x) using seqtk (v1.2-r94) (https://github.com/lh3/seqtk) and a different seed for each down-sampling process to create different fastq files with varying levels of coverage from the original data.

Data Processing

Sequences were trimmed using trimmomatic v0.36 (3), aligned to the respective reference genome with BWA mem v0.7.17 with the minimum seed length parameter set to 100000000 for reproducibility and soft clipping for supplementary alignments (2), and duplicate reads were marked using GATK MarkDuplicatesSpark v4.1.7.0 (4).

Variant Calling

Variants were called in each replicate, each with seven different tools, using multiple parameter configurations for each tool (**Table S1)**. A VCF file containing the intersection of the two replicates was generated using bcftools isec (v1.9) (2). Individual replicate, and intersection VCF files were compared against their respective AF-specific golden VCF using bcftools isec followed by a custom script to parse the results. The pipeline used for data simulation, sequence processing, variant calling, and analysis is available at https://github.com/gencorefacility/MAD2. Variant calls made by each tool were analyzed using custom scripts, available at (https://github.com/GhedinSGS/Optimized-Quantification-of-Intrahost-Viral-Diversity).

Synthetic RNA generation, library preparation, and data processing

Synthetic, 'wildtype' (WT), influenza genomic RNA samples were designed using PB2, HA, and NA segments from the H1N1pdm consensus reference outlined above. Variant RNA was created by adding 18, 14 and 14 known nucleotide changes into the WT PB2, HA, and NA segments respectively. The six genome segments (3 WT and 3 variant) were synthesized as double stranded DNA (gBlocks) and subsequently amplified to incorporate universal influenza A/B virus primer sequences and T7 promoter sequences. The DNA was then *in vitro*-transcribed with the HiScribeTM T7 High Yield RNA Synthesis Kit (Invitrogen) to produce negative sense viral RNA and the concentration of each sample was determined by Qubit. RNA samples were diluted to an equal copy number concentration of $6x10^8$ copies/µL. WT and variant RNA pools were constructed by mixing the three segments at equal molarity. The two RNA pools were mixed at various frequencies (50%, 25%, 12.5%, 6.25%, 3.13%, 1.56%, 0.78%, 0.39%), and diluted to various copy number concentrations ($6x10^6$ - $6x10^3$ copies/µL).

cDNA was generated using Multiplex Reverse Transcription-PCR (Superscript III one step with HiFi platinum *Taq*) for all samples. All libraries were prepared using the Nextera XT library preparation kit (Nextera), scaled down to 0.25x of the manufacturer's instructions, cleaned with AMPure beads, and pooled at equal molarity. Libraries were sequenced on the Miseq 300 Cycle v2 using 2 x 75 pair-end reads. Samples were amplified and sequenced in duplicate and analyzed with the pipeline described above, with the addition of adapter trimming. Synthetic SARS-CoV-2 data from a similarly designed study was downloaded from SRA (PRJNA682212) and processed as above (5).

Sequence analysis of synthetic influenza genome mixtures identified a set of mutations that were reproducibly detected in multiple independent mixtures but were not included in the gBlock design. As we deemed it extremely unlikely that these were recurrent errors introduced by reverse transcription in multiple independent samples, we determined that these variants were likely introduced prior to reverse transcription either during gBlock synthesis, the PCR-mediated introduction of the T7 promoter sequence, or *in vitro* transcription. Thus, these variants were defined as pre-mRTPCR errors and excluded from all analyses.

SARS-CoV-2 clinical sample preparation and sequencing

Total RNA was extracted from 300µL of nasopharyngeal (NP) or mid-turbinate (MT) swabs collected at the NIH Clinical Center as part of diagnostic testing between 07/24/2020 and 03/31/2021 (**Table S2**). All samples were de-identified and anonymized.

Samples were collected and stored in viral transport media (BD, 220220) and RNA was extracted using the QIAamp® Viral RNA Mini Kit (Qiagen, 52904) according to the manufacturer's instructions. Quantitative real-time PCR was performed according to the "CDC Real-Time RT-PCR Panel for Detection 2019-Novel Coronavirus" protocol with three SARS-CoV-2 virus-specific primers/probe sets (N1, N2, N3, (Integrated DNA Technologies, cat. 10006606)) to test for the presence of SARS-CoV-2. A standard curve was generated using the CDC Positive Template Control (PTC) RNA and was used to calculate viral copies/mL.

Amplification of the viral genome was performed using a modified version of the ARTIC consortium protocol for nCoV-2019 sequencing (https://artic.network/ncov-2019) and the methods described at https://github.com/GhedinSGS/SARS-CoV-2_analysis. All libraries were prepared using the Nextera XT library preparation kit (Nextera), scaled down to 0.25x of the manufacturer's instructions. Briefly, PCR products were normalized to 0.2ng/µL. DNA was then fragmented, tagged, amplified, and barcoded (Illumina Nextera DNA dual indexes), cleaned with a 0.9x bead cleanup and pooled at equal molarity. A 0.7x bead cleanup was performed on the final pool and libraries were sequenced on either the Illumina MiSeq or the Illumnia NextSeq500 using either the 2x150 bp or 2x300 bp paired end protocol. All samples were processed in duplicate.

SARS-CoV-2 sequence processing and variant calling

Illumina sequencing adapters and primer sequences were trimmed with Trimmomatic v0.36 (3). Trimmed reads were aligned to the Wuhan-Hu-1 SARS-CoV-2 reference genome (NC_045512.2) using BWA mem v0.7.17 with the -K parameter set to 100000000 for reproducibility and -Y to use soft clipping for supplementary alignments (2). The two resulting SAM files (from A and B primer pools) for each biological sample were merged into one alignment file using Picard Tools MergeSamFiles v2.17.11. Duplicates were marked using GATK MarkDuplicatesSpark v4.1.3.0 (4). Variants were called with each of the 7 variant calling tools, using the above described MAD2 pipeline, with the standard parameters as shown in **Table S1**. All analysis files are available at https://github.com/GhedinSGS/Optimized-Quantification-of-Intrahost-Viral-Diversity .

Vivaldi Package

To facilitate standardized downstream analysis of viral variant calling data, we developed a publicly available R package capable of extracting data from VCF files, calculating common metrics, and generating useful overview plots (https://github.com/GreshamLab/vivaldi). Vivaldi (Viral Variant Location and Diversity) can serve as a starting point for analysis of both simulated and experimental sequencing data.

Synthetic Influenza Viral Sequences

DNA sequences used as templates for in vitro transcription to generate full length synthetic negative sense influenza genome segments.

### HA VARIANT

#### [T7 PROMOTER/Uni13/HA 5' non-coding/HA VAR CDS/HA 3' non-coding/Uni12Inf3]

**TAATACGACTCACTATAGGGAGTAGAAACAAGGGTGTTTTTCCTCATATTTCTGAAATTCTAATC**TTAAATACATATTCTACACT
GTAGAGACCAATTAGAGCACATCCAGAAACTGATTGCCCCCAGGGAGACTACCAGTACCAATGAACTGGCGACAGTTGAATAG
ATCGCCAAAATCTGGTAAATCCTTGTTGATTCCAGTTTTACCCCATCTATTTCTTCTCTGTTTAATTTTGCTTCCTCTGAGTATTTT
GGGTAGTCATAAGTCCCATTTTTGACACTTTCCATGCACGTGTTATCGCATTTGTGGTAAAATTCAAAGCAGCCGTTTCCAATTTC
CTTGGCATTGTTTTTTAGCTGGCTTCTTACCTTTTCATATAAGTTCTTCACATTTGAATCGTGGTAGTCCAAAGTTCTTTCATTTTC
CAATAGAACCAACAGTTCGGCATTGTAAGTCCAAATGTCCAGGAAACCATCATCAACTTTTTTATTTAAATTCTCTATTCTTTTTTC
CAGGTGGTTGAACTCTTTACCTACTGCTGTGAACTGTGTATTCATCTTTTCAATAACAGAATTTACTTTGTTAGTAATTTCGTCAAT
GGCATTCTGTGTGCTCTTCAGGTCGGCTGCATATCCTGACCCCTGCTCATTTTGATGGTGATAACCGTACCATCCATCTACCATC
CCTGTCCACCCCCCTTCAATGAAACCGGCAATGGCCCCAAATAGGCCTCTAGATTGAATAGACGGGATATTCCTCAATCCTGTG
GCCAGTCTCAATTTTGTGCTTTTTACATATTTTGGACATTTTCCAATTGTGATCGGATGTATATTATGAAATGGGAGGCTGGTGTT
TATAGCACCCTTGGGTGTCTGACAAGTTGTATTGCAATCGTGGACTGGTGTATCTGAAATGATAATACCAGATCCAGCGTTTCTT
TCCATTGCGAATGCATATCTCGGTACCAATAGATTTCCAGTTGCTTCGAATGTTATTTTGTCTCCCGGCTCTACTAGTGTCCAGT
AATAGTTCATTCTCCCTTCTTGATCCCTCACTTTGGGTCTTATTGCTATTTCCGGCTTGAACTTCTTGCTGTATCTTGATGTCCCC
ACAAAAACATATGCATCTGCATTCTGATAGAGGCTTTGTTGGTCAGCACTAGTAGATGGATGGTGAATGCCCCATAGCACGAGG
ACTTCTTTCCCTTTATCATTAATGTAGGATTTGCTGAGCTTTGGGTATGAATTCTCTTTTTTAACTAGCCATATTAAATTTTTGTAG
AAGCTTTTTGCTCCAGCATGAGGACATGCTGCCGTTACACCTTTGTTCGAGTCATGATTGGGCCATGAACTTGTCTTGGGGAAT
ATCTCAAACCTTTCAAATGATGACACTGAGCTCAATTGCTCTCTTAGCTCCTCATAATCGATGAAGTCTCCTGGGTAACACGTTC
CATTGTCTGAACTAGGTGTTTCCACAATGTAGGACCATGAGCTTGCTGTGGAGAGTGATTCACACTCTGGATTTCCCAGGATCC
AGCCAGCAATGTTACATTTACCCAAATGCAATGGGGCTACCCCTCTTAGTTTGCATAGTTTCCCGTTATGCTTGTCTTCTAGAAG
GTTAACAGAGTGTGTTACTGTTACATTCTTTTCTAGTACTGTGTCTACAGTGTCTGTTGAATTGTTCGCATGATAACCTATACATA
ATGTGTCTGCATTTGCGGTTGCAAATGTATATAGCAGAACTACTAGTATTGCCTCCAT**TTTGGTTGTTTTTATTTTCCCCTGCTTT
TGCT**

## HA WT

#### [T7 PROMOTER/Uni13/HA 5' non-coding/HA WT CDS/HA 3' non-coding/Uni12Inf3]

**TAATACGACTCACTATAGGGAGTAGAAACAAGGGTGTTTTTCCTCATATTTCTGAAATTCTAATC**TTAAATACATATTCTACACT
GTAGAGACCCATTAGAGCACATCCAGAAACTGATTGCCCCCAGGGAGACTACCAGTACCAATGAACTGGCGACAGTTGAATAG
ATCGCCAAAATCTGGTAAATCCTTGTTGATTCCAGCTTTACCCCATCTATTTCTTCTCTGTTTAATTTTGCTTCCTCTGAGTATTTT
GGGTAGTCATAAGTCCCATTTTTGACACTTTCCATGCACGTGTTATCGCATTTGTGGTAAAATTCAAAGCAGCCGTTTCCAATTTC
CTTGGCATTGTTTTTTAACTGGCTTCTTACCTTTTCATATAAGTTCTTCACATTTGAATCGTGGTAGTCCAAAGTTCTTTCATTTTC
CAATAGAACCAACAGTTCGGCATTGTAAGTCCAAATGTCCAGGAAACCATCATCAACTTTTTTATTTAAATTCTCTATTCTTTTTTC
CAGGTGGTTGAACTCTTTACCTACTGCTGTGAACTGTGTATTCATCTTTTCAATAACAGAATTTACTTTGTTAGTAATCTCGTCAAT
GGCATTCTGTGTGCTCTTCAGGTCGGCTGCATATCCTGACCCCTGCTCATTTTGATGGTGATAACCGTACCATCCATCTACCATC
CCTGTCCACCCCCCTTCAATGAAACCGGCAATGGCCCCAAATAGGCCTCTAGATTGAATAGACGGGACATTCCTCAATCCTGTG
GCCAGTCTCAATTTTGTGCTTTTTACATATTTTGGACATTTTCCAATTGTGATCGGATGTATATTCTGAAATGGGAGGCTGGTGTT
TATAGCACCCTTGGGTGTCTGACAAGTTGTATTGCAATCGTGGACTGGTGTATCTGAAATGATAATACCAGATCCAGCATTTCTT
TCCATTGCGAATGCATATCTCGGTACCACTAGATTTCCAGTTGCTTCGAATGTTATTTTGTCTCCCGGCTCTACTAGTGTCCAGT
AATAGTTCATTCTCCCTTCTTGATCCCTCACTTTGGGTCTTATTGCTATTTCCGGCTTGAACTTCTTGCTGTATCTTGATGTCCCC
ACAAAAACATATGCATCTGCATTCTGATAGAGACTTTGTTGGTCAGCACTAGTAGATGGATGGTGAATGCCCCATAGCACGAGG
ACTTCTTTCCCTTTATCATTAATGTAGGATTTGCTGAGCTTTGGGTATGAATTTCCTTTTTTAACTAGCCATATTAAATTTTTGTAG
AAGCTTTTTGCTCCAGCATGAGGACATGCTGCCGTTACACCTTTGTTCGAGTCATGATTGGGCCATGAACTTGTCTTGGGGAAT
ATCTCAAACCTTTCAAATGATGACACTGAGCTCAATTGCTCTCTTAGCTCCTCATAATCGATGAAATCTCCTGGGTAACACGTTC
CATTGTCTGAACTAGATGTTTCCACAATGTAGGACCATGAGCTTGCTGTGGAGAGTGATTCACACTCTGGATTTCCCAGGATCC
AGCCAGCAATGTTACATTTACCCAAATGCAATGGGGCTACCCCTCTTAGTTTGCATAGTTTCCCGTTATGCTTGTCTTCTAGAAG
GTTAACAGAGTGTGTTACTGTTACATTCTTTTCTAGTACTGTGTCTACAGTGTCTGTTGAATTGTTCGCATGATAACCTATACATA
ATGTGTCTGCATTTGCGGTTGCAAATGTATATAGCAGAACTACTAGTATTGCCTTCAT**TTTGGTTGTTTTTATTTTCCCCTGCTTTT
GCT**

### NA VARIANT

#### [T7 PROMOTER/Uni13/NA 5' non-coding/NA VAR CDS/NA 3' non-coding/Uni12Inf1]

**TAATACGACTCACTATAGGGAGTAGAAACAAGGAGTTTTTTGAACAGA**TTACTTGTCAATGGTAAATGGCAACTCAGCACCGT
CCGGCCAAGACCAACCCACAGTGTCACTGTTTACACCACAAAAAGATATGCTGCTCCCGCTAGTCCAGATTGTGTTCTCTTTGG
GTCGCCCTCTGATTAGTTCAACCCAGAAGCAAGGTCTTATACAATCCAGCCCTGTTAGTTCTGGATGCTGGACAAAACTCCCGC
TATATCCTGACCATTCATTTATTCCTACGATATCTTGCTTTATTGAGAAGTTATTGTCTGTCCCAGTCCATCCGTTCGGATCCCAA

ATCATCTCAAAACCGTTCCTTGAACTAATGCTTTTAGTTCTCCCTATCCAAACACCATTGCCGTATTTGAATGAAAATCCTTTTACT
CCATTTGCTCCATTAGACGATACTGGACCACAACTGCCTGTCTTATCATTAGGGCGTGGATTGTCTCCGAAAATCCCACTACATA
TGTATCCTATCTGATATTCCAGATTCTGGTTGAAAGACACCCACGGTCGATTCGAGCCATGCCAGTTATCCCTGCATACACATGT
GATTTCACTAGAATCAGGATAACAGGAGCATTCCTCATAGTGATAATTAGGGGCATTCATTTCGACTGATTTGACTATCTTTCCCT
TTTCTATTCTGAAGATCTTGTATGAGGCCTGTCCATCACTTGGTCCATCGGTCATTACAGTAAAGCAAGAACCATTTACACATGC
ACATTCAGACTCTTGTGTTCTCAATATATTGTTTCTCCAACTCTTGATAGTGTCTGTTATTATTCCGTTGTACTTTAACACAGCCAC
TGCCCCATTGTCTGGGCCAGAAATTCCAATTGTTAGCCAATTGATGCCATCATGACAAGCACTTGCTGACCAAGCGACTGACTC
AAATCTTGAGTTGTATGGAGAGGGCACTTCACCAATAGGACAGCTCATTAGGGTTCGATATGGGCTCCTGTCTTTAATGGTTCC
ATTGGAATGTTTGTCATTTAGCAAGGCCCCTTGAGTCAAGAAGAAGGTTCTGCATTCCAAGGGGGAGCATGATATGAATGGTTC
CCTTATGACAAACACATCCCCCTTGGAACCGATTCTTACACTGTTGTCTTTACTGTATATAGCCCATCCATCTACAGGGCAGAGA
GAGGAATTGCCCGCTAATTTCACGGAAACCACTGACTGTCCAGCAGCAAAGTTGGTGTTGCTGATGTTAACATATGTCTGATTTA
CCCAAGTGTTGTTTTCATAAGTAATGACGCTTTGATTGCATGTTTCAATCTGATTTTGATTCCCAAGTTGAATTGAGTGGCTAATC
CATATTGAGATTATGTTTCCAATTTGTAATATTAAGTTAGCCATTCCAATTGTCATGCAGACCGAACCAATGGTTATTATCTTTTGG
TTTGGATTCAT**TTTAAACC****CCTGCTTTCGCT**

**NA WT**

**[T7 PROMOTER/Uni13/NA 5’ non-coding/NA WT CDS/NA 3’ non-coding/Uni12Inf1]**

**TAATACGACTCACTATAGGGAGTAGAAACAAGGAGTTTTTTGAACAGA**TTACTTGTCAATGGTAAATGGCAACTCAGCACCGT
CTGGCCAAGACCAACCCACAGTGTCACTGTTTACACCACAAAAGGATATGCTGCTCCCGCTAGTCCAGATTGTGTTCTCTTTGG
GTCGCCCTCTGATTAGTTCAACCCAGAAGCAAGGTCTTATACAATCCAGCCCTGTTAGTTCTGGATGCTGAACAAAACTCCCGC
TATATCCTGACCACTCATTTATTCCTACGATATCTTGCTTTATTGAGAAGTTATTGTCTGTCCCAGTCCATCCGTTCGGATCCCAA
ATCATCTCAAAACCGTTTCTTGAACTAATGCTTTTAGTTCTCCCTATCCAAACACCATTGCCGTATTTGAATGAAAATCCTTTTACT
CCATTTGCTCCATTAGACGATACTGGACCACAACTGCCTGTCTTATCATTAGGGCGTGGATTGTCTCCGAAAATCCCACTGCATA
TGTATCCTATCTGATATTCCAGATTCTGGTTGAAAGACACCCACGGTCGATTCGAGCCATGCCAGTTATCCCTGCACACACATGT
GATTTCACTAGAATCAGGATAACAGGAGCATTCCTCATAGTGATAATTAGGGGCATTCATTTCGACTGATTTGACTATCTTTCCCT
TTTCTATTCTGAAGATCTTGTATGAGGCCTGTCCATCACTTGGTCCATCGGTCATTACAGTAAAGCAAGAACCATTTACACATGC
ACATTCAGACTCTTGTGTTCTCAATATATTGTTTCTCCAACTCTTGATAGTGTCTGTTATTATGCCGTTGTACTTTAACACAGCCAC
TGCCCCATTGTCTGGGCCAGAAATTCCAATTGTTAGCCAATTGATGCCATCATGACAAGCACTTGCTGACCAAGCGACTGACTC
AAATCTTGAGTTGTATGGAGAGGGAACTTCACCAATAGGACAGCTCATTAGGGTTCGATATGGGCTCCTGTCTTTAATGGTTCC
ATTGGAATGTTTGTCATTTAGCAAGGCCCCTTGAGTCAAGAAGAAGGTTCTGCATTCCAAGGGGGAGCATGATATGAATGGTTC
CCTTATGACAAACACATCCCCCTTGGAACCGATTCTTATACTGTTGTCTTTACTGTATATAGCCCATCCACTAACAGGGCAGAGA
GAGGAATTGCCCGCTAATTTCACGGAAACCACTGACTGTCCAGCAGCAAAGTTGGTGTTGCTGATGTTAACATATGTCTGATTTA
CCCAAGTGTTGTTTTCATAAGTAATGACGCTTTGATTGCATGTTTCAATCTGATTTTGATTCCCAAGTTGAATTGAGTGGCTAATC
CATATTGAGATTATGTTTCCAATTTGTAATATTAAGTTAGCCATTCCAATTGTCATACAGACCGAACCAATGGTTATTATCTTTTGG
TTTGGATTCAT**TTTAAACC****CCTGCTTTCGCT**

**PB2 VARIANT**

**[T7 PROMOTER/Uni13/PB2 5’ non-coding/PB2 VAR CDS/PB2 3’ non-coding/Uni12Inf1]**

**TAATACGACTCACTATAGGGAGTAGAAACAAGGTCGTTTTTAAACTATTCGACA**ATTAATTGATGGCCGCCCGAATTCTTTTGG
TCGCTGTCTGGCTGTCAGTAAGTATGCTAGAGTCCCGTTTTCGTTTCATTACCAACACTACGTCCCCTTGCCCAATTAGCACATT
AGCGTTCTCTCCTTTTGCAAGATTGCTCAGTTCATTGATGCTTAATGCTGGGCCATATCTCTTGTCTTCTTTGCCCAAAATGAGAA
ATCCTCTCAGGACAGCAGACTCCACTCCAGATGTGCCTTCATCTGGATCTTCAGTCAATGCACCTGCATCCTTTCCAAGAACTGT
AAGTCGTTTGGTTGCCTTGTTGTAATTGAATACTGGAGAATTGCCTCTTACCAGTATCCTCAACCCTGATCCTCTCACATTCACA
GTCAATGAGGAGAATTGCATCCTACTCTGTTCTGGTGGAGCAGCAGCAAAGGGGAGAAGTTTTATTATTTGGACAGTGTCAAAT
GTCCCAAGCACATCCCGCATTTGCTGGAACAGTGTCCTTACAAATCCACTGTACCGGCTTCTGGTTGCCTTAGGGACAAGAGAC
TGAAATGGTTCAAATTCCATTTTGTTGTATAACATTGTGGGATCTTGTGACCATTGAATTTTCACAATTTCCCAGTTCCTGATTATC
CATTGATAAGTGTTGACTAGCACTGACTCAGGGCCATTGATCTCCCACATCATTGATGACGAATAAGTTATTGTCAACCTCTCAG
TTCCTTGCGTTTCACTGACTTCTTCGGGAGACAATAGTACGTTCCCTCTTTGATCTCTAACCCTTAAAAATCGGTCAATACTCACT
ACCACTCTCTCCGTGCTGGAGTATTCATCTACTCCCATTTTGCTGACTCTTATCCCTCTCAGCGACATCTCCATGCTTGGGGTCA
TGTCGGGCAGTATTCCGATCATTCCCATCACATTGTCGATGGATTCAATTCCCCAGTTCTGGAAAAGCACTTTTGCATCTTTTTG
GAAATGCCTCAAGAGTTGGTGCATTGGGTTCAGTCGCTGGTTTGCCCTATTGACAAAGTTCAGATCGCCCCTAACTGCCTTGAT
CATGCAATCCTCTTGTGAGAATACCATGGCCACAATTATTGCCTCAGCAATTGACTGCTCGTCCCTCCCGCTTACTATCAACTGG
ATCAATCTCCTGGTTGCCTTTCTGAGAATAGCTGTTGCTCTTCTCCCAACCATTGTGAATTCTTCATACCCTTCATGTACTCTTAT
TTTCAGTGTTTGGAGGTTGCCCGTTAGCACTTCTTCTTCATTCTTGACTGATGATCCGCTTGTCCTTTTGAAAGTGAACCCACCA
AAACTGAAAGATGAGCTAATCCTCAACCCTATTGCTGCCTTGCATATGTCTACGGCTTGTTCCTCAGTTGGATTCTGTCTAAGGA
TATCCACCATCCTTACTCCTCCAATCTGTGTGCTGTGGCACATTTCCAAGAGAGATGCTAATGGGTCTGCTGACACTGCTGCTCT
TCTTACTATGTTTCTAGCAGCGATAATCAAACTTTGGTCAACATCATCATTTCTCACTTCTCCTCCTGGAGTGTACATCTGCTCCC
AGCACGTTCCTTGGGTTAAGTGCAACACTTCAATATAAACACTGCCTGTTCCGCCGGCTACTGGGAGAAACCTTGTTTTACGGA
CCAATTCTCTTTCTAGCATGTACGCCACCATCAAGGGAGCAATTTTACAATCCTGGAGCTCTTCTTTCTTCTCTTTTGTTATTGCC
AGTTGTGACTCTGATGTCAGTATTCTTGCCCCCACTTCATTTGGGAAAACAACTTCCATAATCACATCCTGTGCCTCCTTGGCAC
TGAGATCTGCATGGCCAGGGTTTGTATCAACTCTCCTCCTTATTTTAACTTGATTTCTGAAGTGGACAGGGCCGAAGGTACCATA
TTTCAACCTTTCGACCTTTTCGAAATAAGTTTTATATACCTTAGGGTAATGAACTGTACTTGTTGTTGGGCCATTCCTATTCCACC

ATGTTACGGCCAGAGGTGATACCATCACTCGGTCTGATCCAGCATCGTTTGTTTTGCTCCAGAGGGTTTGCCCTTGTTCATTCCT
CTCTGGAATCATGTCCATTATTCTCTTGTCTGCTGTAATTGGGTATCTCATTGCCATCATCCACTTCATTCTGAGTGCGGGGTTCT
TTTCTTGCCTTCCTGATGTGTACTTTTTGATTATGGCCATATGGTCCACAGTGGTCTTAGTGAGTATCTCGCGAGTGCGGGACTG
CGACATTAGATCTCTCAGTTCTTTTATTCTCTCCAT**ATTGAATATAATTGA**CCTGCTTTCGCT

**PB2 WT**

**[T7 PROMOTER/Uni13/PB2 5' non-coding /PB2 WT CDS/PB2 3' non-coding/Uni12Inf1]**

**TAATACGACTCACTATAGGGAGTAGAAACAAGGTCGTTTTTAAACTATTCGACA**ACTAATTGATGGCCATCCGAATTCTTTTGG
TCGCTGTCTGGCTGTCAGTAAGTATGCTAGAGTCCCGTTTTCGTTTCATTACCAACACTACGTCCCCTTGCCCAATTAGCACATT
AGCGTTCTCTCCTTTTGCAAGATTGCTCAGTTCATTGATGCTTAATGCTGGGCCATATCTCTTGTCTTCTTTGCCCAAAATGAGAA
ATCCTCTCAGGACAGCAGACTCCACCCCAGATGTGCCTTCATCTGGATCTTCAGTCAATGCACCTGCATCCTTTCCAAGAACTG
TAAGTCGTTTGGTTGCCTTGTTGTAATTGAATACTGGAGAATTGCCTCTTACCAGTATCCTCAACCCTGATCCTCTCACATTCACA
GTCAATGAGGAAAATTGCATCCTACTCTGTTCTGGTGGAGCAGCAGCAAAGGGGAGAAGTTTTATTATTTGGACAGTGTCAAAT
GTCCCAAGCACATCCCGCATTTGCTGGAACAGTGTCCTTACGAATCCACTGTACCGGCTTCTGGTTGCCTTAGGGACAAGAGA
CTGAAATGGTTCAAATTCCATTTTGTTGTATAACATTGTGGGATCTTGTGACCATTGAATTTTCACAATTTCCCAGTTCCTGATTAT
CCATTGATAAGTGTTGACTAGCACTGACTCAGGGCCATTGATCTCCCACATCATTGATGACGAATAAGTTATTGTCAACTTCTCA
GTTCCTTGCGTTTCACTGACTTCTTCGGGAGACAATAGTACGTTCCCTCTTTGATCTCTAACCCTTAAAAATCGGTCAATACTCAC
TACCACTCTCTCCGTGCTGGAGTATTCATCTACTCCCATTTTGCTGACTCTTATCCCTCTCAGCGACATCTCCGTGCTTGGGGTC
ATGTCGGGCAGTATTCCGATCATTCCCATCACATTGTCGATGGATTCAATTCCCCAGTTCTGGAAAAGCACTTTTGCATCTTTTT
GGAAATGCCTCAAGAGTTGGTGCATGGGGTTCAGTCGCTGGTTTGCCCTATTGACAAAGTTCAGATCGCCCCTAACTGCCTTGA
TCATGCAATCCTCTTGTGAGAATACCATGGCCACAATTATTGCCTCAGCAATTGACTGCTCGTCTCTCCCGCTTACTATCAACTG
GATCAATCTCCTGGTTGCCTTTCTGAGAATAGCTGTTGCTCTTCTCCCAACCATTGTGAATTCTTCATACCCTTCATGTACTCTTA
TTTTCAGTGTTTGGAGGTTGCCCGTTAGCACTTCTTCTTCTTTCTTGACTGATGATCCGCTTGTCCTTTTGAAAGTGAACCCACCA
AAACTGAAAGATGAGCTAATCCTCAACCCTATTGCTGCCTTGCATATGTCTACGGCTTGTTCCTCAGTTGGATTCTGTCTAAGGA
TGTCCACCATCCTTACTCCTCCAATCTGTGTGCTGTGGCACATTTCCAAGAGAGATGCTAATGGGTCTGCTGACACTGCTGCTC
TTCTTACTATGTTTCTAGCAGCGATAATCAAACTTTGGTCAACATCATCATTTCTCACTTCTCCTCCTGGAGTGTACATCTGCTCC
CAGCACGTCCCTTGGGTTAAGTGCAACACTTCAATATAAACACTGCCTGTTCCGCCGGCTACTGGGAGAAACCTTGTTTTACGG
ACCAATTCTCTTTCTAGCATGTACGCCACCATCAAGGGAGCAATTTTACAATCCTGGAGCTCTTCTTTCTTCTCTTTTGTTATTGC
CAGCTGTGACTCTGATGTCAGTATTCTTGCCCCCACTTCATTTGGGAAAACAACTTCCATAATCACATCCTGTGCCTCCTTGGCA
CTGAGATCTGCATGGCCAGGGTTTGTATCAACTCTCCTCCTTATTTTAACTTGATTTCTGAAGTGGACAGGGCCGAAGGTACCAT
GTTTCAACCTTTCGACCTTTTCGAAATAAGTTTTATATACCTTAGGGTAATGAACTGTACTTGTTGTTGGGCCATTCCTATTCCAC
CATGTTACGGCCAGAGGTGATACCATCACTCGGTCTGATCCAGCATCGTTTGTTTTGCTCCAGAGGGTTTGTCCTTGTTCATTCC
TCTCTGGAATCATGTCCATTATTCTCTTGTCTGCTGTAATTGGGTATCTCATTGCCATCATCCACTTCATTCTGAGTGCGGGGTTC
TTCTCTTGCCTTCCTGATGTGTACTTTTTGATTATGGCCATATGGTCCACAGTGGTCTTAGTGAGTATCTCGCGAGTGCGGGACT
GCGACATTAGATCTCTCAGTTCTTTTATTCTCTCCAT**ATTGAATATAATTGA**CCTGCTTTCGCT


REFERENCES

1. Wybo WA, Jordan J, Ellenberger B, Marti Mengual U, Nevian T, Senn W. 2021. Data-driven reduction of dendritic morphologies with preserved dendro-somatic responses. Elife 10.

2. Li H, Durbin R. 2009. Fast and accurate short read alignment with Burrows-Wheeler transform. Bioinformatics 25:1754-60.

3. Bolger AM, Lohse M, Usadel B. 2014. Trimmomatic: a flexible trimmer for Illumina sequence data. Bioinformatics 30:2114-20.

4. Van der Auwera GA, Carneiro MO, Hartl C, Poplin R, Del Angel G, Levy-Moonshine A, Jordan T, Shakir K, Roazen D, Thibault J, Banks E, Garimella KV, Altshuler D, Gabriel S,

DePristo MA. 2013. From FastQ data to high confidence variant calls: the Genome Analysis Toolkit best practices pipeline. Curr Protoc Bioinformatics 43:11 10 1-11 10 33.

5. Valesano ALR, Kalee E; Dimcheff, Derek E; Blair, Christopher N; Fitzsimmons, William J; Petrie, Joshua G; Martin, Emily T; Lauring, Adam S. 2021. Temporal dynamics of SARS-CoV-2 mutation accumulation within and across infected hosts. bioRxiv doi:https://doi.org/10.1101/2021.01.19.427330.
